# Inositol phosphorlyceramide synthase null *Leishmania major* are viable and virulent in animal infections where salvage of host sphingomyelin predominates

**DOI:** 10.1101/2022.06.14.496188

**Authors:** F. Matthew Kuhlmann, Phillip N. Key, Suzanne M. Hickerson, John Turk, Fong-Fu Hsu, Stephen M. Beverley

**Affiliations:** Department of Molecular Microbiology, Washington University School of Medicine, 660 S. Euclid Ave., Saint Louis, MO 63110, USA; Department of Internal Medicine, Washington University School of Medicine, 660 S. Euclid Ave., Saint Louis, MO 63110, USA

**Keywords:** Trypanosomatid protozoan parasite, Sphingolipids, inositol phosphorylceramide, lipid remodeling, lipid salvage, infectivity and virulence

## Abstract

Many pathogens synthesize inositolphosphorylceramide (IPC) as the major sphingolipid (SL), differing from the mammalian host where sphingomyelin (SM) or more complex SLs predominate, and the divergence between IPCS and mammalian sphingolipid synthases has prompted interest as a potential drug target. However, in the trypanosomatid protozoan *Leishmania*, cultured insect stage promastigotes lacking *de novo* sphingolipid synthesis (Δ*spt2^-^*) and sphingolipids entirely survive and remain virulent, as infective amastigotes salvage host sphingolipids and continue to produce IPC. To further understand the role of IPC, we generated null IPCS mutants in *L. major* (Δ*ipcs^-^*). Unexpectedly and unlike fungi where IPCS is essential, Δ*ipcs^-^* was remarkably normal in culture and highly virulent in mouse infections. Both IPCS activity and IPC were absent in Δ*ipcs^-^* promastigotes and amastigotes, arguing against an alternative route of IPC synthesis. Notably, salvaged mammalian sphingomyelin (SM) was highly abundant in purified amastigotes from both WT and Δ*ipcs^-^*, and salvaged SLs could be further metabolized into IPC. SM was about 7-fold more abundant than IPC in WT amastigotes, establishing that SM is the dominant amastigote SL, thereby rendering IPC partially redundant. These data suggest that SM salvage likely plays key roles in the survival and virulence of both WT and Δ*ipcs^-^* parasites in the infected host, confirmation of which will require the development of methods or mutants deficient in host SL/SM uptake in the future. Our findings call into question the suitability of IPCS as a target for chemotherapy, instead suggesting that approaches targeting SM/SL uptake or catabolism may warrant further emphasis.

## Introduction

Leishmaniasis is often considered a neglected tropical disease. Currently it is estimated that there are more than 1.7 billion people at risk and nearly 12 million people with symptomatic disease, ranging from mild cutaneous infections to severe disfiguring or lethal forms, with upwards of 50,000 deaths annually (1–4). Remarkably, the great majority of infections are undiagnosed and/or asymptomatic, suggesting that well more than 100 million people may harbor parasites (3, 5). Persistent asymptomatic infections constitute a double edged sword – while serving as a reservoir for transmission and/or reactivation by stress or immunosuppression, persistent parasites are known to mediate strong protective concomitant immunity against disease pathology (3,5–7). *Leishmania* have two distinct growth stages, a promastigote stage in the sand fly vector, and an intracellular amastigote stage residing within cellular endocytic pathways in the mammalian host (4). While the promastigote stage is readily cultured in the laboratory and amenable to molecular techniques, amastigotes require the use of macrophage infection systems or infections of animal models replicating key aspects of human disease.

While progress has been achieved in new chemotherapeutic strategies, many are not yet implemented clinically and resistance to several promising agents has already been observed (8–10). One pathway potentially offering important drug target is that of sphingolipid synthesis (11–13). Sphingolipids (SL) are a diverse class of lipids that function in apoptosis, cell signaling, and membrane structure and these pathways have been targeted for therapies against a wide range of diseases including cancer, multiple sclerosis, and infectious diseases (14–18). Sphingolipids differ in the nature of the head groups as well as the composition of the hydrophobic ceramide anchor, in different tissues and species. Unlike mammalian cells that primarily synthesize sphingomyelin (SM) and complex glycosphingolipids, *Leishmania* synthesize inositol phosphorylceramide (IPC) as their primary sphingolipid (19), as do other protozoa, fungi and plants (11,20,21).

This divergence has prompted scrutiny of IPCS as a drug target, where in fungi it is essential and repression causes decreased growth and virulence along with increased susceptibility to acidic environments (22, 23). IPC is formed when inositol phosphorylceramide synthase (IPCS) converts ceramide and phophatidylinositol (PI) to IPC and diacylglycerol (DAG). The related trypanosomatid parasite *Trypanosoma brucei* encodes four sphingolipid synthases (SLS), collectively mediating synthesis of sphingomyelin (SM), ethanolamine phosphorylceramide and IPC; inducible RNAi knockdowns of the entire locus were lethal, although the contribution of the individual genes/activities was not further dissected (24). *Leishmania* IPCS has been pursued as a drug target by analogy of findings from pathogenic fungi as well as its divergence from mammalian sphingomyelin synthases (25). Aureobasidin A, a known inhibitor of yeast IPCS, inhibits parasite growth at high concentrations, probably off-target from IPCS inhibition (25), but other candidate inhibitors have been advanced (26–30).

Notably, the biological requirement for *Leishmania* IPCS has not been addressed *in vivo*. That *Leishmania* could differ from other organisms was first suggested by the fact that *L. major* null mutants lacking the first enzyme in *de novo* sphingolipid synthesis, serine palmitoyl transferase (Δ*spt2^-^*) completely lacked all SLs, yet remain viable in log phase culture (31, 32). Subsequent studies in promastigotes showed that sphingolipid catabolism was used to generate ethanolamine (EtN), whose provision to Δ*spt2^-^* parasites rescued a defect in differentiation to infective metacyclic parasites (33). While Δ*spt2^-^* was fully infective in animal models, it continued to synthesize IPC, presumably by salvage of potential precursor sphingolipid components from the host cell (34–36). Several routes could be envisaged; sphingoid (long-chain) bases or ceramide precursors could be salvaged directly, or mammalian sphingolipids could be acquired and catabolized for resynthesis as IPC sphingolipids. As all routes ultimately require IPCS for parasite IPC synthesis, we sought to generate Δ*ipcs^-^* null mutants if possible, and to explore their impact on IPC synthesis and/or metabolism, and virulence in animal models. Surprisingly, our data show convincingly *IPCS* was not essential, with SM salvage in WT cells rendering it the dominant amastigote SL as well as serving as the source of amastigote IPC. These findings likely account for the dispensability of IPCS in animal infections, potentially dampening enthusiasm for IPCS as a useful drug target in this parasite.

## Results

### Targeted Replacement of *L. major IPCS*

The *L. major* genome is primarily diploid albeit with occasional aneuploidy, and typically two rounds of gene replacement are required to generate null mutants of single copy genes (37). Targeting constructs containing resistance markers to G418 (*NEO)* or nourseothricin (*SAT*) were designed to successively replace the *IPCS* ORF precisely by homologous gene replacement (Fig. 1A). To alleviate concerns about lethality of IPCS deficiency, we focused on the culturable promastigote stage where SLs are not required in any capacity in the presence of ethanolamine (31–33). A secondary effect of IPCS deficiency could be ceramide accumulation which can be toxic (38), and we also performed transfections in the presence of myriocin (an inhibitor of *de novo* SL synthesis) to potentially ameliorate this.

**Figure 1:**
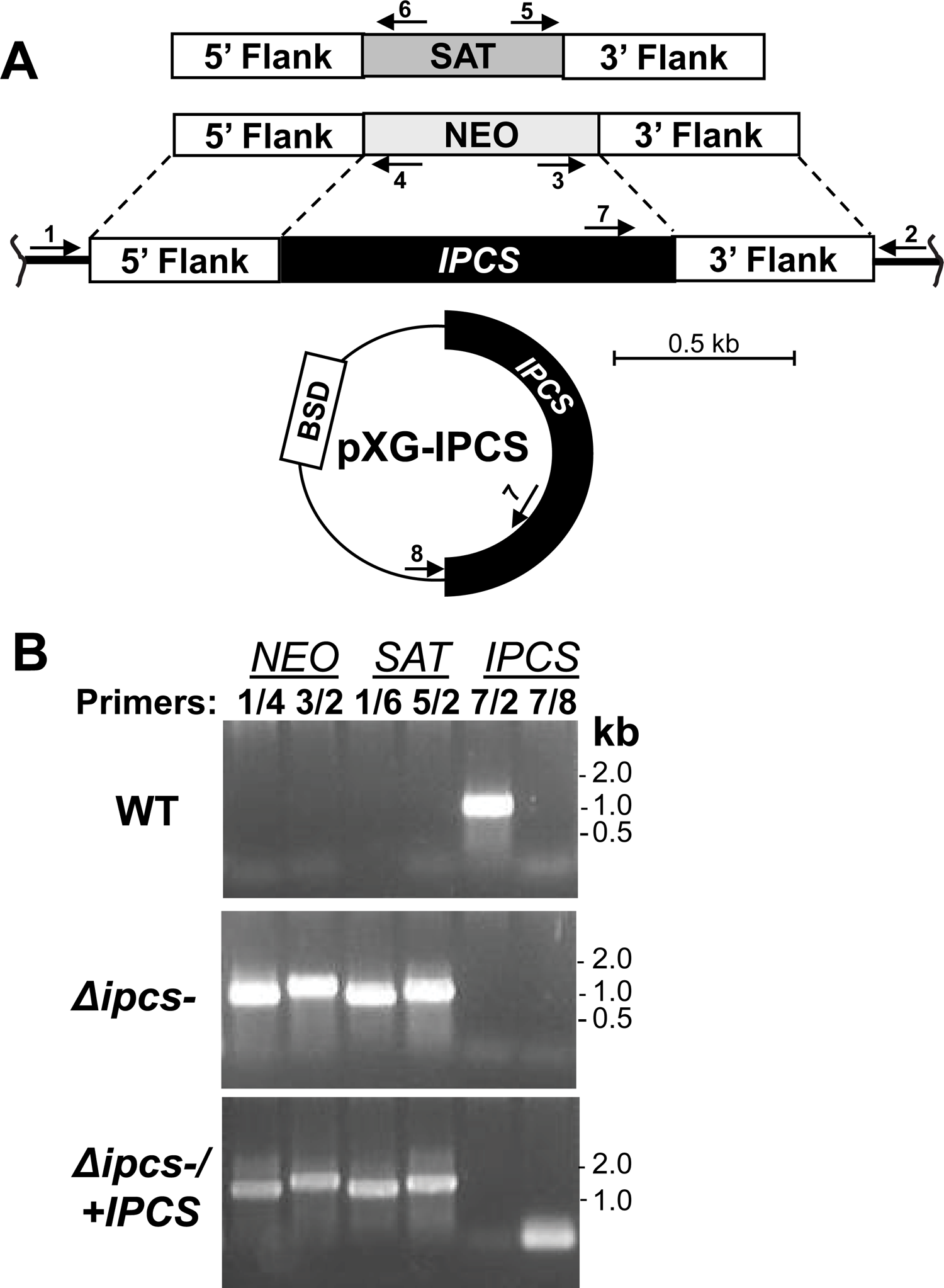
Generation of *Δipcs*^-^ promastigotes. (A) Schematic of planned replacements for *IPCS* with drug markers for nourseothricin (*SAT*), G418 (*NEO*) and expression construct pXG-IPCS with blasticidin (*BSD*) resistance. Arrows represent the location of primers used for PCR; Primers 1/4 and 3/2 tests for correct *NEO* 5’and 3’ replacements; primers 1/6 and 5/2 test for correct *SAT* 5’ and 3’ replacement; primers 7/2 tests for WT *IPCS* and 7/8 test for the pXG-born *IPCS*. Primer 1, SMB 2598; 2, SMB 2827; 3, SMB 2600; 4, SMB 2608; 5, SMB 2609; 6, SMB 2601; 7, SMB 2607; and 8, SMB 1421; sequences can be found in Table S1. (B) PCR confirmation of parasite transfectants. *IPCS*::*NEO* replacement (lanes 1,2, primers 1+4, expected size 860bp; and 3+2, expected size 1024bp); planned *IPCS::SAT* replacement (lanes 3,4, primers 1+6, expected size 849bp; and 5+2, expected size 933bp); WT *IPCS* (lane 5, primers 7+2, expected size 882bp); and pXG-BS-IPCS (lane 6, primers 7+8, expected size 124bp).

Heterozygous (*IPCS/IPCS::ΔNEO*) and homozygous null mutants (*IPCS*::Δ*SAT*/*IPCS*::Δ*NEO*; referred to as *Δipcs***^-^**) were readily obtained and confirmed by PCR tests to lack the IPCS coding region, accompanied by the expected planned replacements (analysis of the double homozygous replacement is shown in Fig. 1B). An episomal expression vector (pXG-BSD-IPCS) was used to restore *IPCS* expression, yielding the line *Δipcs*^-^*/+IPCS* (Fig. 1 A,B). The recovery of *Δipcs***^-^** clonal lines was readily accomplished with good efficiencies, regardless of EtN or myriocin treatment, and these showed no strong growth defects. Numerous transfectant lines were obtained and authenticated with similar phenotypes, and in this work results from one representative set of KO and IPCS-restored pair are shown.

### IPCS activity is lost in *Δipcs*^-^ promastigotes

Enzymatic assays with promastigote microsomes were performed by evaluating the conversion of phosphatidylinositol (PI) and NBD-Ceramide to NBD-IPC, after TLC separation, and quantitative fluoroscopy (Figs. 2A, S1A). High levels of NBD-IPC were generated in complete reactions bearing microsomes, NBD-Cer and PI, and this product was sensitive to PI-PLC digestion as expected. We purified and confirmed the presumptive NBD-IPC from the TLC plate by mass spectrometry^1^. Consistent with prior studies, addition of 20 µM aureobasidin A showed little inhibition of the *in vitro* reaction (26). Little or no product was obtained if PI or NBD-Cer were omitted, if PC replaced PI, or if microsomes were omitted or previously heat inactivated. These data confirmed our ability to assay IPCS, with the predicted enzymatic properties (Figs. 2A, S2A).

**Figure 2:**
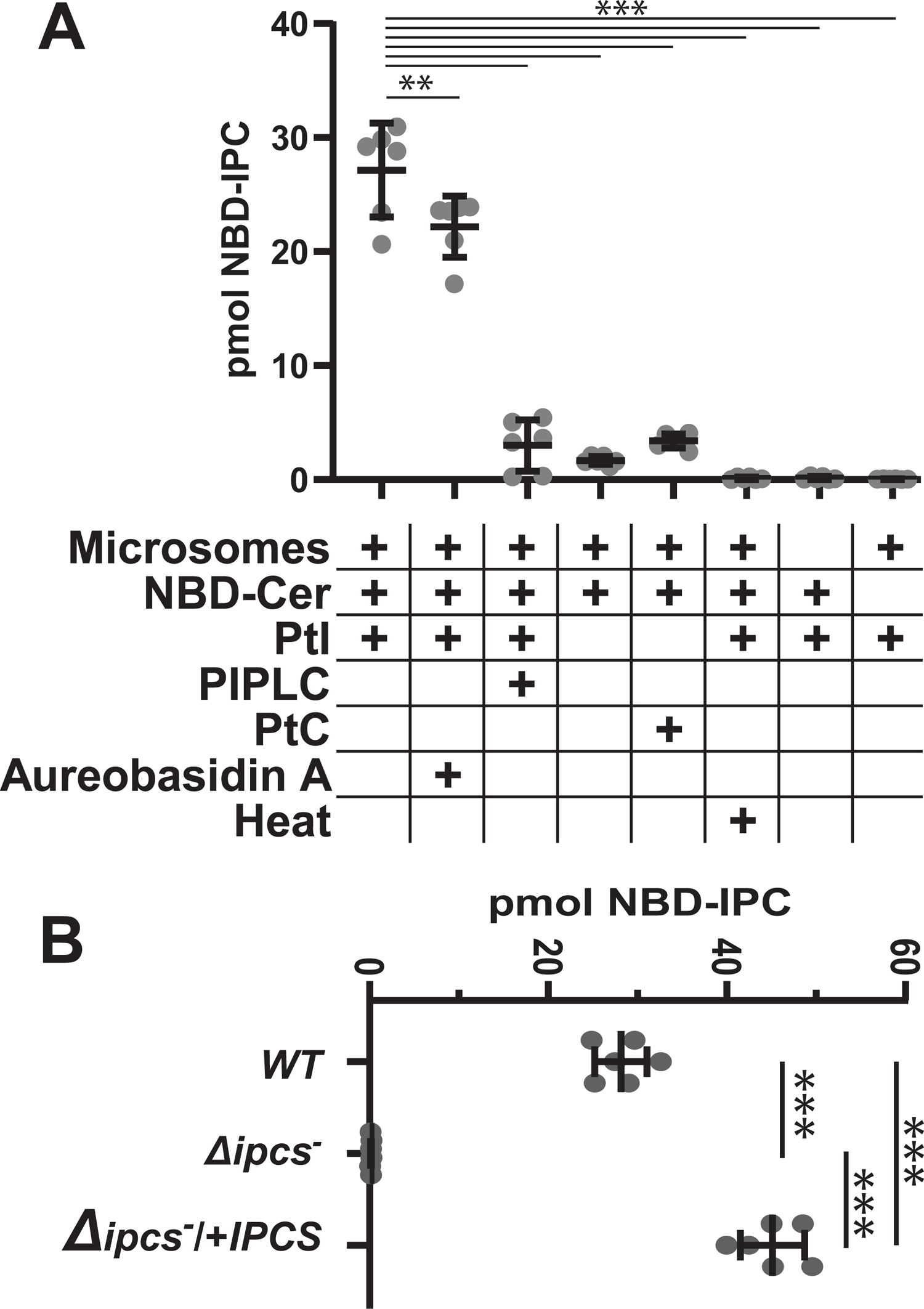
Enzymatic characterization of IPCS activity and its absence in Δ*ipcs^-^* promastigotes. (A) NBD-IPC formation in complete reactions or in the absence of key substrates, microsomes, inhibitors (20 µM aureobasidin A), or enzymatic treatments, as indicated. Statistical comparisons to the standard reaction (column 1) are shown; **, *p* <002; ***, *p*< .0001 (ANOVA with Dunnet’s post-hoc analysis). (B) Quantitation of NBD-IPC formation by microsomes from WT, Δ*ipcs^-^* or Δ*ipcs^-^*/+*IPCS* parasites. ***, *p* < .0001 (ANOVA with Tukey’s post-hoc analysis). For both panels, primary TLC data can be found in Fig. S1, 3 experiments with 2 technical replicates were performed, and bars represent the mean ± 1 SD.

We compared the IPCS activity of WT, *Δipcs***^-^**/+ or *Δipcs***^-^**/+ *IPCS* parasite microsomes (Figs. 2B, S1B). As expected, *Δipcs***^-^** showed no detectable activity, while *Δipcs***^-^**/+ IPCS showed about 50% more than WT (*p* < 0.001).

### *Δipcs*^-^ promastigotes lack IPC

Mass spectrometry was used to evaluate lipids present in whole cell extracts of WT, *Δipcs***^-^** and *Δipcs***^-^***/+IPCS* promastigotes (Fig. 3A-C). In WT, two ion peaks at *m/z* 778.6 and 806.6 corresponding to [M – H]^-^ ions of d16:1/18:0- and d18:1/18:0–IPC respectively were seen, as described previously (32,34,39). Consistent with the genetic deletion of *IPCS*, these IPC ions were undetectable in *Δipcs***^-^** promastigotes. IPC levels were restored in the *Δipcs***^-^/+***IPCS* promastigotes, albeit only partially to about 50% WT levels (Fig. 3C).

**Figure 3:**
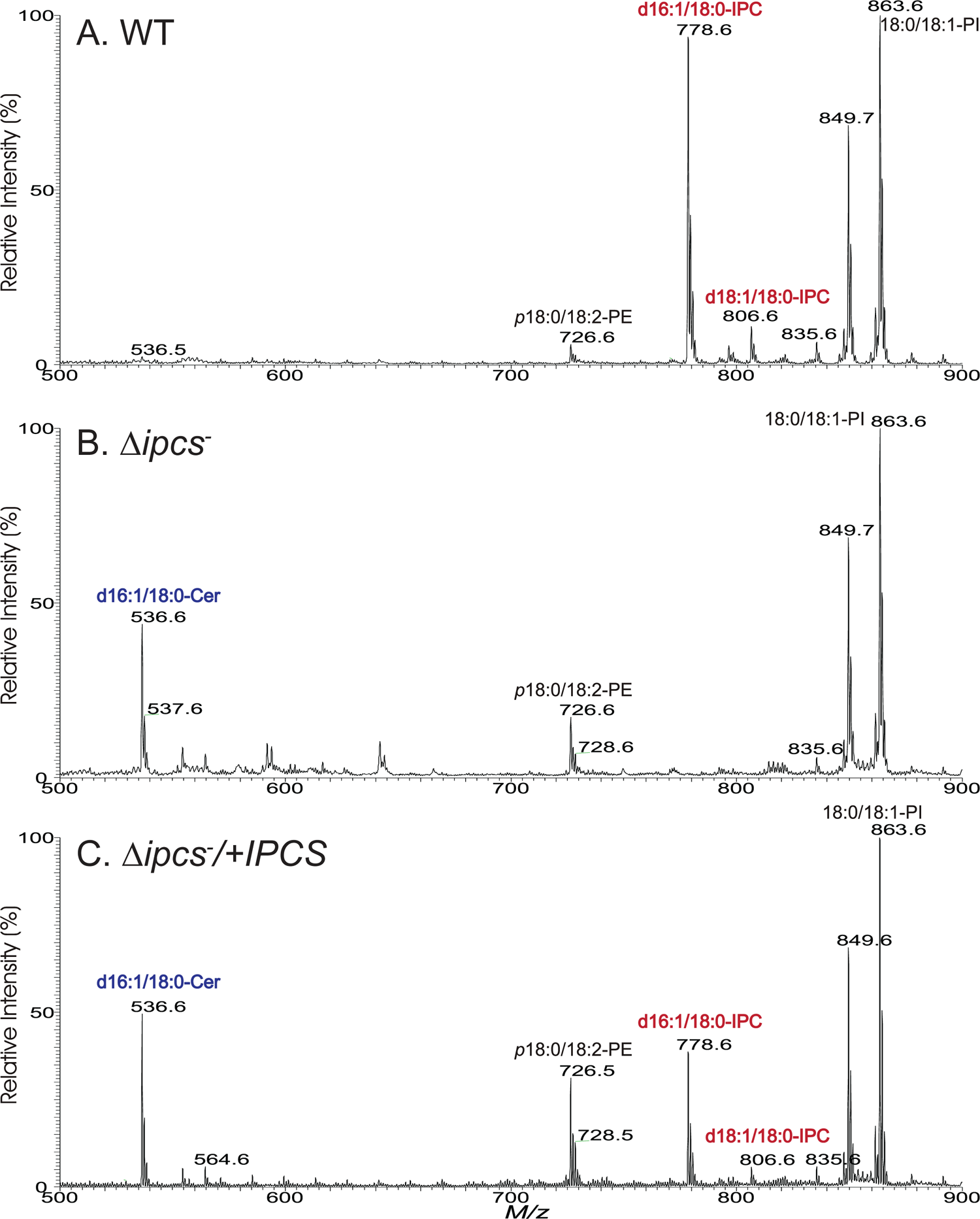
Absence of IPC in Δ*ipcs^-^ Leishmania*. ESI/MS spectra (negative ion mode) obtained with lipid preparations from logarithmic growth phase WT, Δ*ipcs^-^*, or Δ*ipcs^-^*/+*IPCS* promastigotes.

Together with the results from enzymatic assays, these data established that *Leishmania* IPCS is the sole source of IPCS enzymatic activity or IPC.

As predicted, IPC precursor d16:1/18:0-ceramide (*m/z* 536.6) levels were elevated in the *Δipcs***^-^** promastigotes (Fig. 3B) relative to WT cells (Fig. 3A). Curiously, this ceramide peak did not decrease noticeably in *Δipcs***^-^***/+IPCS,* despite restoration of IPC levels (Fig. 3C). Small variations in the levels of PE ions of *m/z* 726.6 (*p*18:0/18:2-PE) and 728.6 (*p*18:0/18:1-PE) were seen, but the levels of other lipids identified by MS such as ions at *m/z* 849 (e18:0/18:1-PI) and 863 (18:0/18:1-PI), did not appear to change in either *Δipcs***^-^** or *Δipcs***^-^***/*+*IPCS* promastigotes (Fig. 3A-3C). The causes of these minor changes were not pursued further.

### *Δipcs*^-^ promastigotes remain viable and form metacyclics

As promastigotes *in vitro, Δipcs***^-^** grew somewhat slower than WT, with a doubling time of 9.0 ± 0.7 vs. 7.7 ± 0.4 hr for WT (mean ± 1 SD, n=3), reaching a stationary phase density of about 70% WT (*p* < 0.05; Fig. 4A). Upon entry into stationary phase a small percentage of nonviable cells emerged, as judged by propidium iodide exclusion (Fig. 4B). Since *Δipcs***^-^** promastigotes should retain sphingolipids capable of generating EtN (33), we suspected that provision of EtN would have no effect on stationary phase viability, which proved correct (Fig. S2 A,B). Restoration of IPCS expression in the *Δipcs***^-^***/+IPCS* line returned growth and viability to WT (Fig. 4A,B).

**Figure 4:**
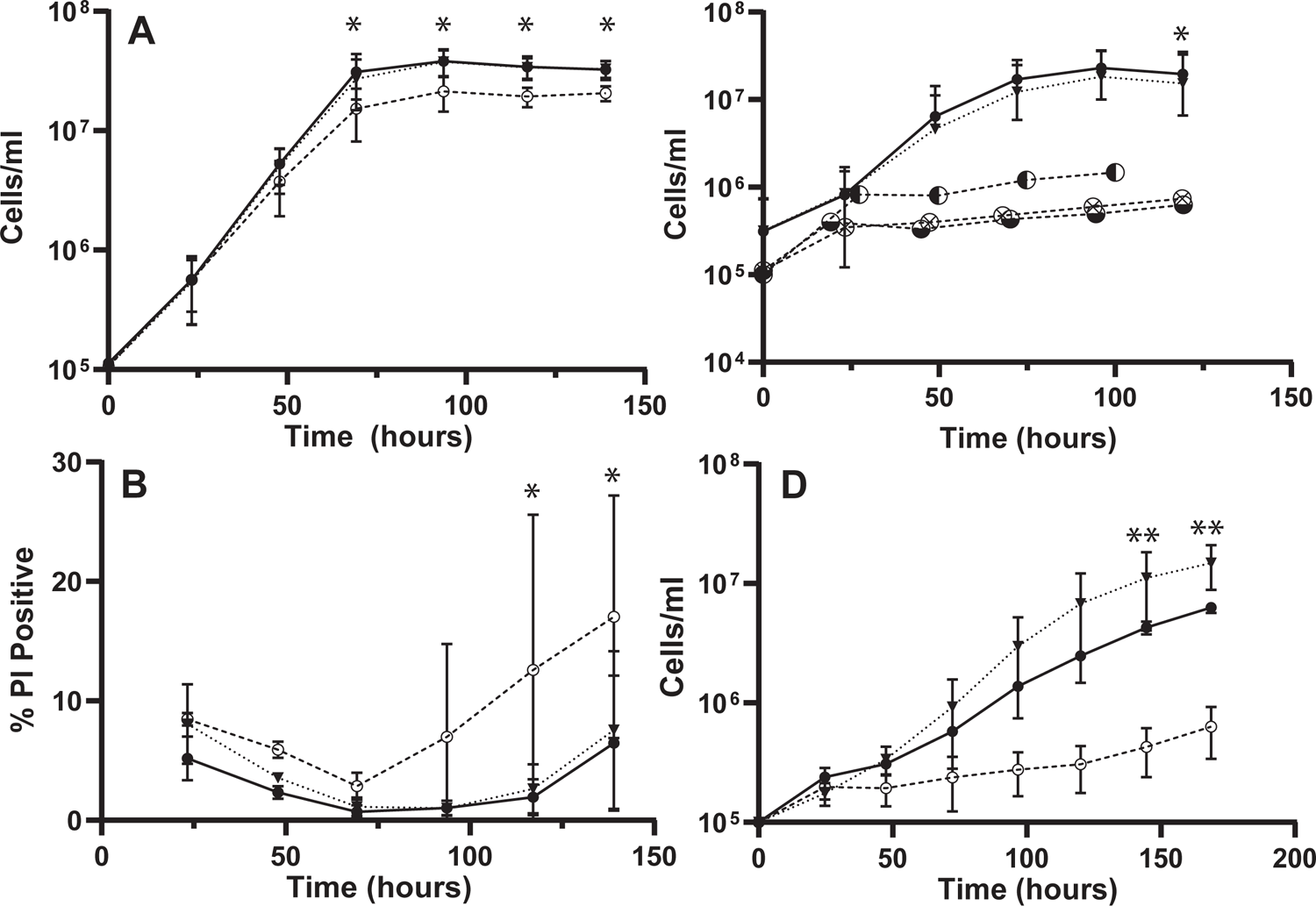
Phenotypic characterization of Δ*ipcs^-^* promastigotes *in vitro*. (A), growth, and (B), Propidium iodide exclusion (viability), 7 experimental replicates (C), growth in the presence of 2 μM 3-keto-dihydrosphingosine (⊗), dihydrosphingosine (◐) or phytosphingosine (◒; n=1 for each) and; (D), growth at pH 5.0 (MES buffer), 2 experimental replicates. All data points are mean ± 1SD, ANOVA tests for each panel were significant (*p* ≤ 0.002) followed by Bonferroni correction to adjust for multiple comparisons, * post-hoc comparisons between WT and Δ*ipcs^-^* significant at p <0.05, ** post-hoc comparisons between Δ*ipcs^-^*/+IPCS and Δ*ipcs^-^* significant at p <0.05. All parasites were propagated in standard M199 culture media at 25⁰ C. Lines tested are WT (●, solid line), Δ*ipcs^-^* (○), or Δ*ipcs^-^*/+IPCS (▾, dashed line).

Upon entry into stationary phase *Leishmania* promastigotes differentiate to an infective metacyclic form that can be purified by lectin or density-gradient methods (40). *Δipcs***^-^** showed a small reduction in metacyclics relative to WT or the *Δipcs***^-^***/+IPCS* control (5.7 ± 2.7 % in *Δipcs***^-^** vs. 11.4 ± 7.1 % in WT, or 10.4 ± 5.1% in *Δipcs***^-^***/+IPCS*; mean ± SD, n = 2; Fig. S2 C; these differences were not statistically significant).

### *Δipcs*^-^ promastigotes are hyper-susceptible to exogenous sphingoid bases

Accumulation of ceramide, sphingoid bases (SBs), or their phosphorylated forms, is toxic to most eukaryotic cells (38). This suggested that in the absence of further metabolism by IPCS, addition of exogenous SBs would likely lead to excess toxic ceramide or SB levels in *Δipcs***^-^**. Accordingly, daily addition of 2 μM 3-keto-dihydrosphingosine, dihydrosphingosine, or C17 phytosphingosine resulted in death of *Δipcs***^-^** parasites (Fig. 4C), which was rescued in *Δipcs***^-^/+***IPCS*. For cultures receiving C17 phytosphingosine, mass spectrometry confirmed synthesis of phytoceramide (Fig. S3).

### *Leishmania Δipcs*^-^ promastigotes are susceptible to extreme pH but not temperature stress

*IPCS* null *Cryptococcus neoformans* is susceptible to pH but not temperature stress (22), and we obtained similar results for *L. major Δipcs*^-^ promastigotes. At pH 4.0 the WT, *Δipcs*^-^ and *Δipcs*^-^/+IPC lines all died, while at pH 5.0, *Δipcs*^-^ grew much slower than WT or *Δipcs*^-^/+IPC (Fig. 4D). At pH 5.5 and above no significant differences were seen amongst the three lines (Supplemental Table S2). Temperature stress was tested at 30° and 33° C with no significant effects on the relative growth of WT, *Δipcs*^-^ and *Δipcs*^-^/+IPC lines (Fig. S2 D); at 37° C, all cells died.

### *Δipcs*^-^ defects are not reversed by diacylglycerol

The IPCS product DAG also functions as a messenger in multiple signaling pathways mediated by Protein Kinase C (PKC). *Leishmania* lack a known PKC although it has been suggested that other kinases may fulfill similar roles (41). We tested the ability of exogenous DAG to rescue the growth or viability reductions seen in *Δipcs*^-^ promastigotes by adding DAG daily to cultures. However, with two separate supplementations (0.2 or 2 μM) no improvement was observed (Figs. S2E,F), suggesting that DAG deficiency did not contribute to *Δipcs*^-^ phenotypes. While we did not confirm DAG uptake by parasites here, that exogenous DAG is accessible to *Leishmania* was shown previously, as a brief 30 min exposure to 250 nM DAGs resulted in stimulation of transferrin endocytosis (41).

### *Δipcs*^-^ promastigotes are rounded but lipid bodies and acidocalcisomes are unchanged

In Δ*spt2^-^* promastigotes, sphingolipid deficiency causes changes in cell rounding, acidocalcisomes, and lipid bodies (33, 34). In *Δipcs*^-^, with a small increase in rounded cells observed (21.6 ± 0.6% vs. 7.4 ± 2.0% for WT or 8.9 ± 3.0% for *Δipcs*-/*+IPCS,* 2 replicates; *p* < 0.02 for comparisons of the mutant vs. the other two lines). WT and *Δipcs*^-^ appeared similar by DAPI staining of acidocalcisomes/polyphosphates, or Nile Red O staining for lipid accumulation and lipid bodies (Fig. S4).

### *Δipcs*^-^ remain fully virulent in mouse infections

To test the key question of whether *IPCS* is essential for virulence, we infected susceptible BALB/c or γ-interferon knockout C57BL/6 mice with 1 x 10^5^ metacyclic promastigotes. Unexpectedly the *Δipcs*^-^ infections progressed significantly faster than those of WT or *Δipcs*^-^*/+IPC,* which were similar (a representative experiment is shown in Fig. 5 and combined data from four experiments is shown in Fig. S5 A). Limiting dilution assays at key time points showed that regardless of genotype, lesions of similar sizes had similar parasite numbers (Fig. S5 B), establishing that increased lesion size reflected increased parasite numbers primarily. These data show that if anything, *Δipcs*^-^ was unexpectedly somewhat hypervirulent.

**Figure 5:**
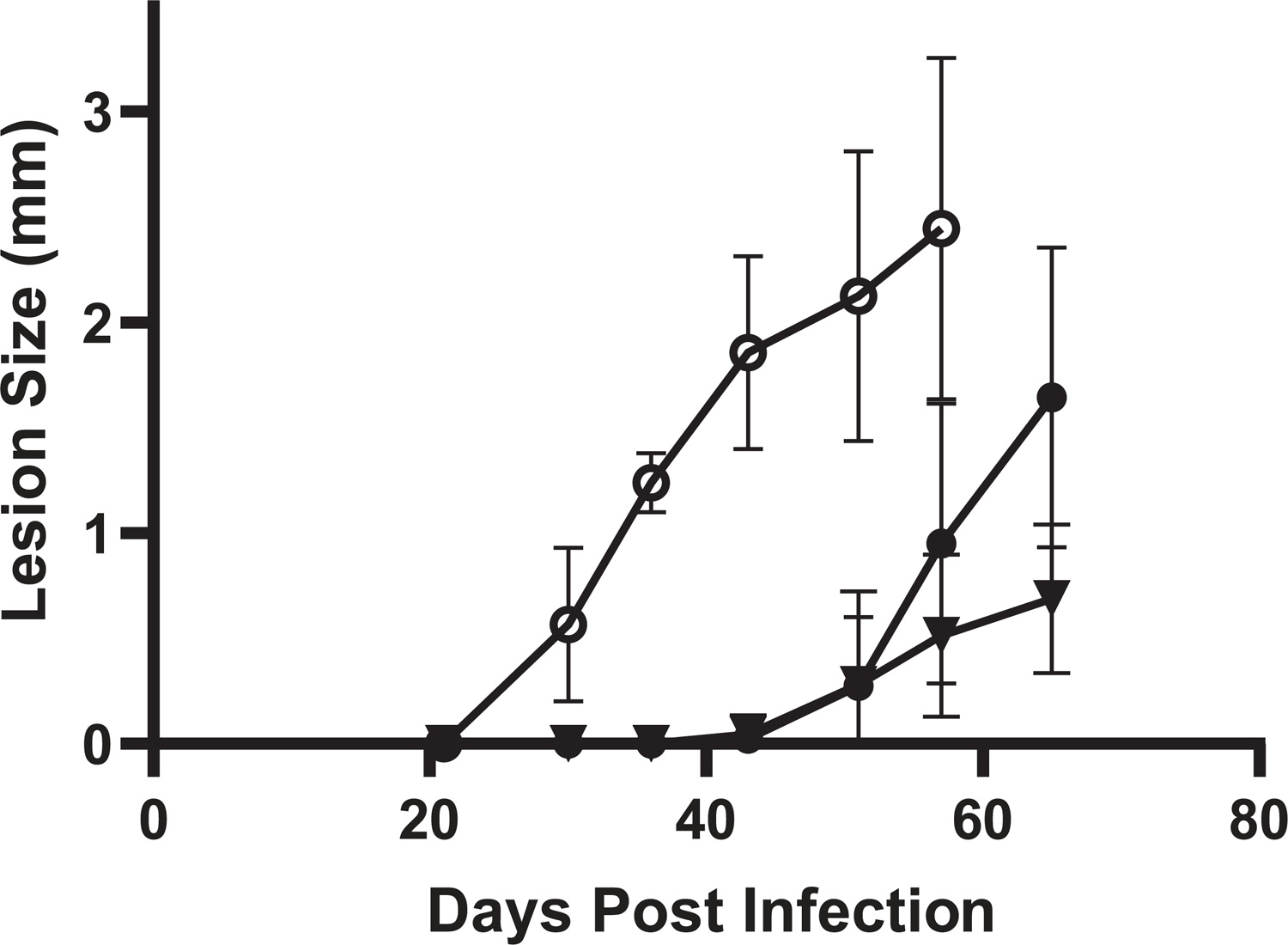
Δ*ipcs^-^* promastigotes are fully virulent in susceptible mice. 10^5^ purified metacyclic parasites were inoculated into the footpad of Balb/C mice and the lesion size measured. Data from a representative experiment is shown, with the average and standard deviation for groups of 4 mice; simple linear regression determined significantly different slopes between groups (p < 0.0001); combined data from 4 separate experiments show similar trends and are provided in Fig. S5 A. Lines tested are WT (●), Δ*ipcs^-^* (○), or Δ*ipcs^-^*/+IPCS (▾).

### *Δipcs*^-^ amastigotes lack IPC

To eliminate the possibility that amastigotes elaborated an alternative IPCS activity, parasites were inoculated into susceptible mice, and purified from similarly sized lesions after 1-2 months. Lipids were then extracted for analysis by electrospray ionization mass spectrometry (ESI/MS) in the negative ion mode. High amounts of background signal complicated interpretations of full scan spectra (Fig. S6), which was overcome by employing linked scan spectra using precursor ion scan of *m/z* 241 specific for detection of [M – H]^-^ ions of PI and IPC lipids (39, 42). These scans showed abundant IPC species (ions of *m/z* 778.6, 806.6 and 808.6) in purified WT amastigotes, as seen previously (Figs. 6A, S7A,G) (34). In *Δipcs*^-^ amastigotes, only trace signals in the relevant *m/z* regions spanning the IPC *m/z* peaks of 778, 806 and 808 were evident, in four separate purified preparations (Figs. 6B, S7C,E,H,I). These peaks were considered background, first by virtue of their low signal/noise ratio (values <3 are considered noise and <10 insufficient for quantitation (43) and secondly, a lack of the expected natural isotope pattern for real peaks (44). In contrast, these criteria validate the assigned IPC peaks in the WT purified amastigotes or infected tissues. The abundance of other lipid species did not appear to change consistently or significantly (Figs. S6-8). No significant differences were seen in amastigotes purified from BALB/c or γ-IFN knockout (γKO) C57BL/6 mice (Figs. S6-8).

**Figure 6:**
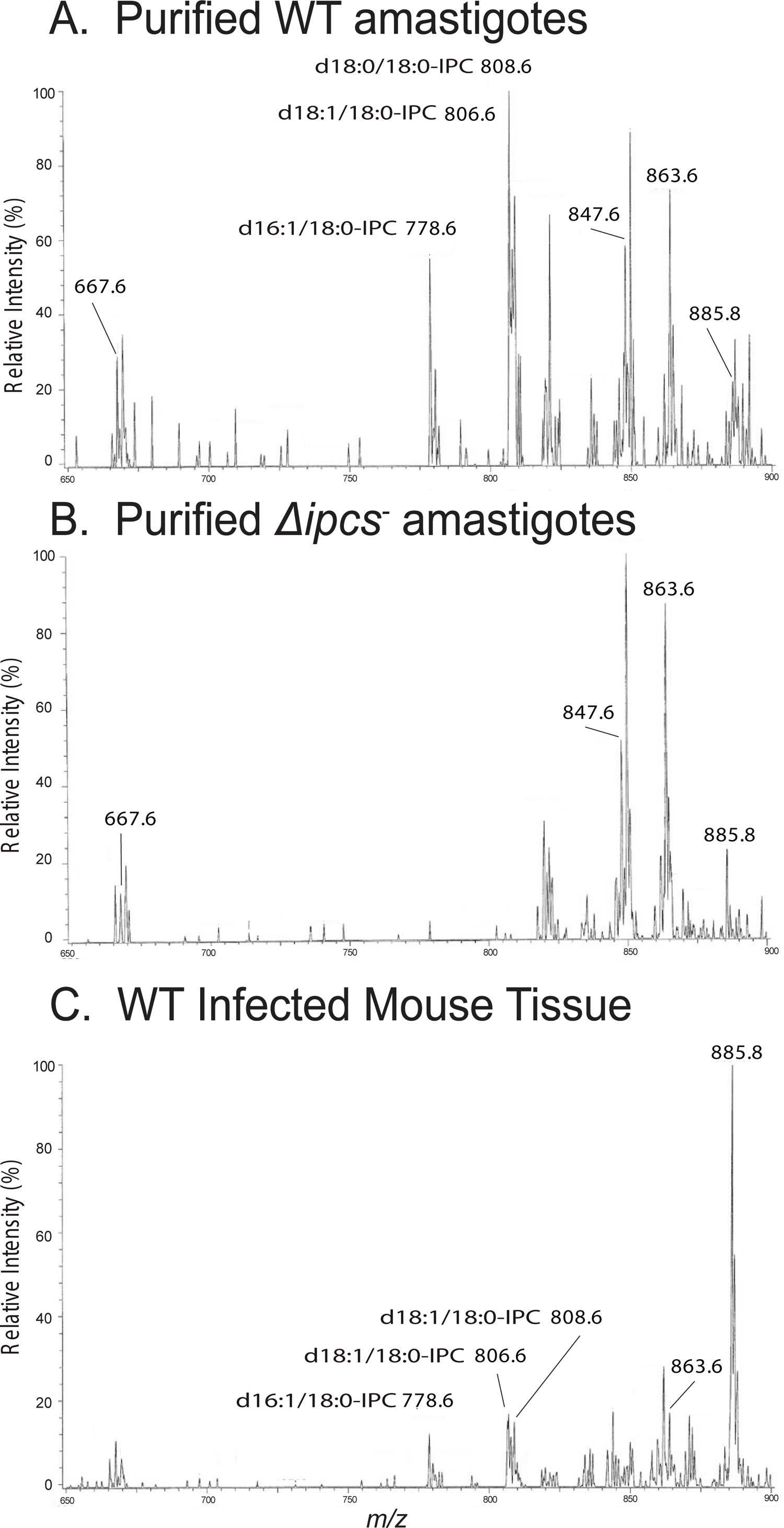
ESI/MS analysis (negative ion mode) of IPCs from purified amastigotes and infected tissue. Spectra were obtained by a triple stage quadrupole instrument applying precursor scan of 241 in the negative ion mode that focuses on IPCs and inositol phospholipid species in the lipid extracts. Samples were prepared from (A) purified WT amastigotes, (B) purified Δ*ipcs^-^* amastigotes (clone WAE), and (C) total infected WT footpad lesions. γKO mice were used in the studies shown; similar findings with BALB/c hosts are shown in Figs. S6 and S7. Abbreviations: d16:1/18:0-IPC, phosphoryl inositol N-stearoylhexadecesphing-4-enine); d18:1/18:0-IPC, phosphoryl inositol N-stearoylsphingosine; d18:0/18:0-IPC, phosphoryl inositol N-stearoylsphinganine.

Thus, purified *Δipcs*^-^ amastigotes lack detectable IPC, and the ability of *Δipcs*^-^ *L. major* to survive and induce pathology is not dependent on its presence.

### WT amastigotes contain abundant host-derived SM

Previous studies showed that *L. major* Δ*spt2*^-^ parasites lacking the key enzyme for *de novo* SB base synthesis nonetheless showed high levels of IPC within infected animals, suggesting that *Leishmania* amastigotes have the ability to salvage host sphingolipids (34, 45). We examined this more directly by MS in purified amastigotes from both WT and *Δipcs*^-^ parasites, which revealed high levels of ions corresponding to SM (Figs. 7A,B, S8). While in many mammalian tissues the most abundant form of SM is d18:1/18:0 (46), in our preparations the most abundant SM was d18:1/16:0 (Figs. 7, S8). This agrees with earlier findings that d18:1/16:0-SM is the most abundant in macrophage preparations commonly used in the study of *Leishmania*, including thioglycolate-elicited peritoneal, bone marrow derived or RAW.264.7 macrophages (47, 48) (Table S3). SM quantitation was achieved by measuring the ratio of the peak height of the [M + Na]^+^ ion of d18:1/16:0-SM (*m/z* 725.7) to that of a spike-in control d18:1/12:0-SM (*m/z* 669.5) (Fig. 7A-C). The data showed that SM in WT infected mouse tissue was found to be about 1.3 times that of the purified *Δipcs*^-^ amastigotes from two preparations (the increase was not statistically significant).

**Figure 7:**
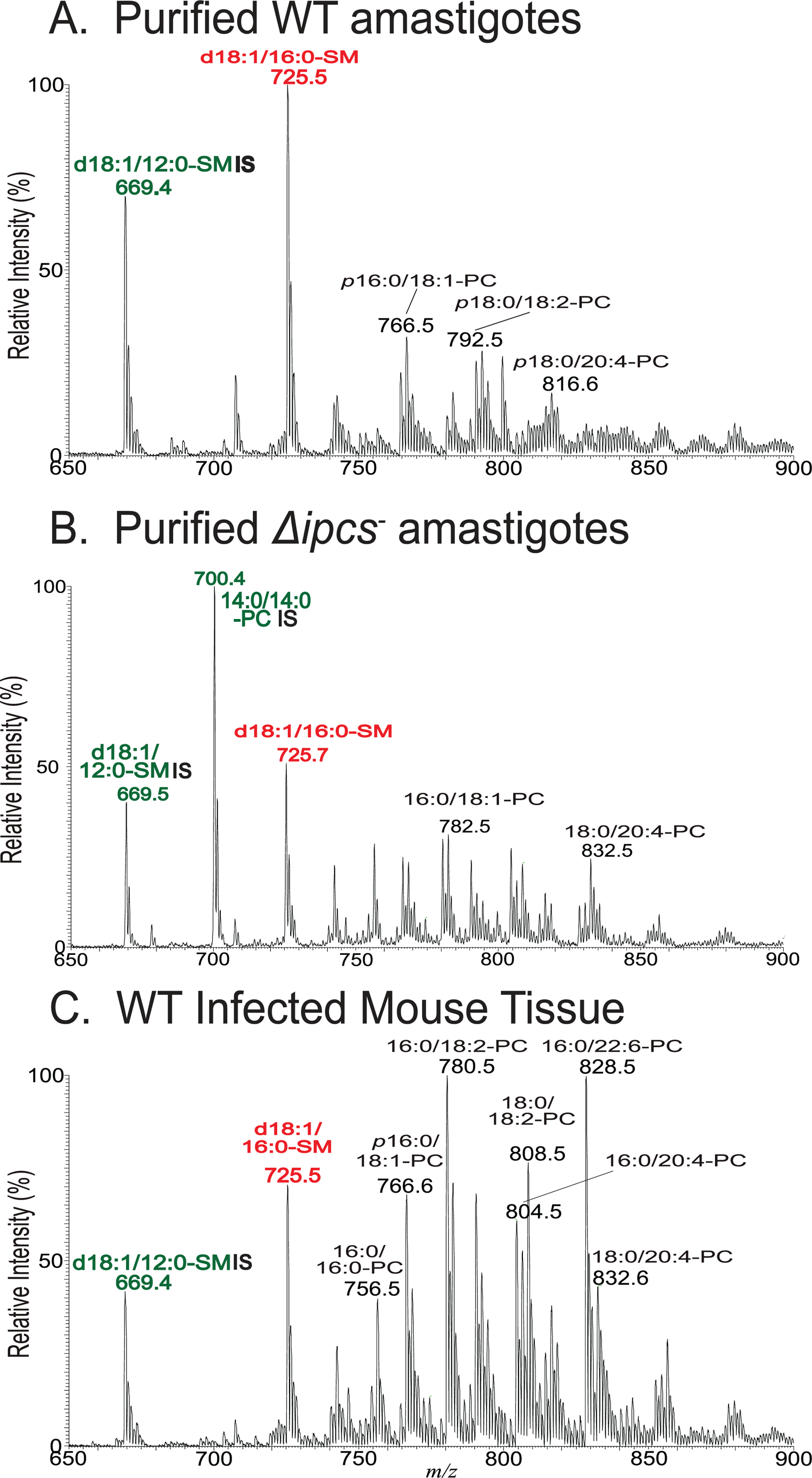
ESI/MS analysis (positive ion mode) of SMs from purified amastigotes and infected tissues. Spectra were acquired in the positive ion mode showing the [M + Na]+ ions of SM (red) and PC, PE or TG (black). Ions labeled in green are spike-in controls (d18:1/12:0 SM or 14:0/14:0-PC). Lipids were prepared from (A) purified WT amastigotes, (B) purified Δ*ipcs^-^* amastigotes (clone WAE), and (C) total WT infected footpad lesions. γKO host mice were used in the studies shown; similar findings with BALB/c hosts are shown in Fig. S8. Abbreviations: d18:1/16:0-SM, N-palmitoyl-octadecesphing-4-enine-1-phosphocholine; PC, phosphatidylcholine; PE, phosphatidylethanolamine; SM, sphingomyelin; TG, triacylglycerol.. For phospholipids (PL) designation: A/B-PL illustrates A: combined number of carbon chain of the fatty acid (FA) attached to glycerol backbone; B: total number of unsaturated bonds of FA substituents.

These data suggested that the levels of salvaged mammalian SM were remarkably high in purified amastigotes. However, purified amastigotes typically are contaminated by host material, which can be readily visualized in lipid extracts from total infected footpad lesions where the mouse tissue contributions dominate (Fig. 7C). We preferred this control instead of uninfected mouse footpads, which lack the extensive infiltration of immune cell types with differing lipid profiles within infected lesions. We sought an estimate of the degree of host contamination of our purified amastigotes, a goal challenged by the complexity of the samples and mix of overlapping and unique lipids between *Leishmania* and mouse. Qualitatively, the host contamination was relatively minor, as it can be seen by the relative decrease of diverse PC species in purified amastigotes vs. the whole WT infected tissue (Fig. 7A,B vs 7C). We estimated a relative loss of mouse lipids to be about 16-fold, based on the major PC species, such as 34:1-, 34:2-, 36:2-, 36:4-, 38:6- and 40:6-PC (Fig. 7; ranging from 9-26-fold). This was consistent with the 9-fold decreased levels of IPC seen in infected tissues relative to purified amastigotes (Fig. 6A,C; calculated relative to the prominent 18:0/20:4-PI peak at *m/z* 885.8).

Notwithstanding some caution that the ion (*m/z* 885) is a suitable internal control, the data clearly suggest that most of the SM in purified amastigote preparations must originate from the parasite itself rather than contaminating host material.

### Salvaged SM and not IPC is the predominant SL in amastigotes

To corroborate the findings above, we evaluated the relative amounts of IPC and SM in WT amastigotes, where IPC is quite abundant (34). We employed a spike-in approach where non-indigenous standards expected to have similar MS ionization behavior were added to each sample, titrating the amounts in pilot studies until a similar peak height was obtained with the biological species of interest (34). The amastigote preparations were spiked with 10 nmol N-Lauryl SM and 2 nmol dipalmitoyl PI, followed by positive or negative ion ESI/MS (Fig. 8). For IPC, the amastigote IPC species peak heights sum to about 90% of that of the PI standard, while the amastigote SM peak height was about 130% of the SM standard (Fig. 8), suggesting the relative amounts of IPC:SM was 1:7 in the purified amastigotes. As noted earlier, only small changes in this value could be attributed to host contamination, and thus, SM is likely the dominant sphingolipid in WT amastigotes. While this assay could not be applied to *Δipcs*^-^ amastigotes, the similar levels of SM in these relative to WT SM must likewise dominate in the mutant (Figs. 7A,B; S8),

**Figure 8:**
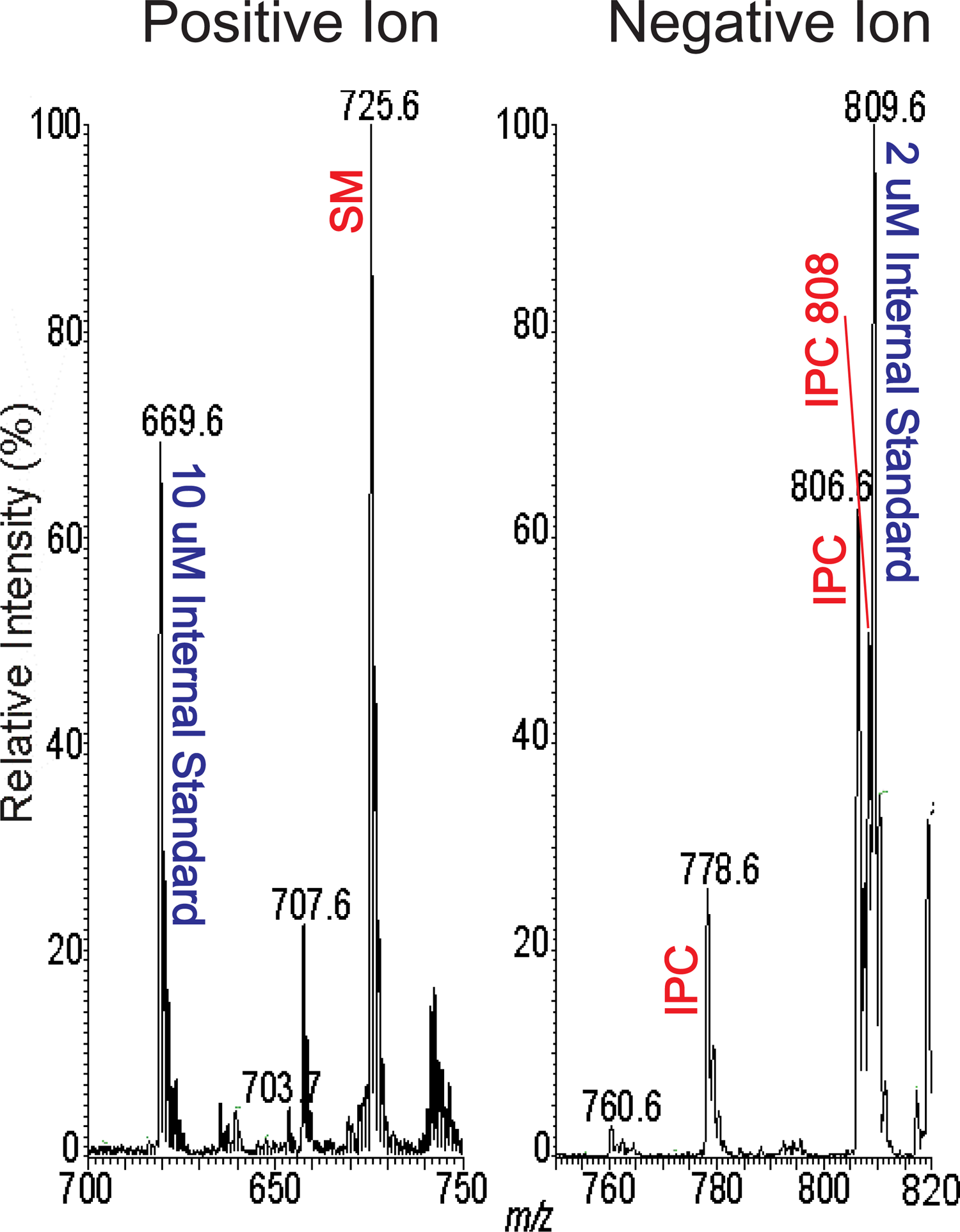
Relative abundance of WT amastigote SM and IPC. Internal standards (10 nmol d18:1/12:0-SM and 2 nmol of dipalmitoyol phosphoinositol) were added to WT amastigote samples, and lipids were extracted and analyzed by ESI/MS in the positive ion mode (panel A) or negative ion mode (panel B) to compare the relative amounts of SM and IPC present in the same sample. This experiment was repeated a second time with similar results.

### WT amastigotes contain IPC with mammalian-like ceramide anchors

The two promastigote IPC ions at *m/z* 778.6 and 806.6 have d16:1/18:0- and d18:1/18:0-Cer backbones, respectively (32,39,47). In contrast, amastigotes showed three IPC species, two of the same *m/z* as in promastigotes but additionally one of m/z of 808.6; the *m/z* 806 and 808 were more abundant in amastigotes and the *m/z* 778 was greater in promastigotes (Fig. 6; (34)). To further explore their origins, we employed ESI/MS to determine the structures.

For the [M – H]^-^ ion of IPC at *m/z* 778.6, the MS^2^ spectra from both amastigotes and promastigotes contain a peak at *m/z* 598 resulting from the loss of inositol (Fig. 9A). MS^3^ on m/z 598 originated from amastigotes (Fig. 9B) yielded ions of *m/z* 360 and 342, arising from loss of 16:0-FA as acid and ketene, respectively, together with ion of *m/z* 255 representing a 16:0 fatty acid carboxylate anion, leading to a final structure of d18:1/16:0-IPC. In contrast, MS^3^ on m/z 598 originated from promastigotes (Fig. 9C) yielded ions of *m/z* 332 and 314 arising from loss of 18:0-FA as acid and ketene, respectively, together with the 18:0 fatty acid carboxylate anion ion at *m/z* 283, resulting in assignment of a d16:1/18:0-IPC structure (32, 39). Interestingly, and unlike promastigotes, the amastigote IPC ion of *m/z* 778.6 contains the same ceramide backbone as the abundant salvaged mammalian SM (Fig. 7).

**Figure 9:**
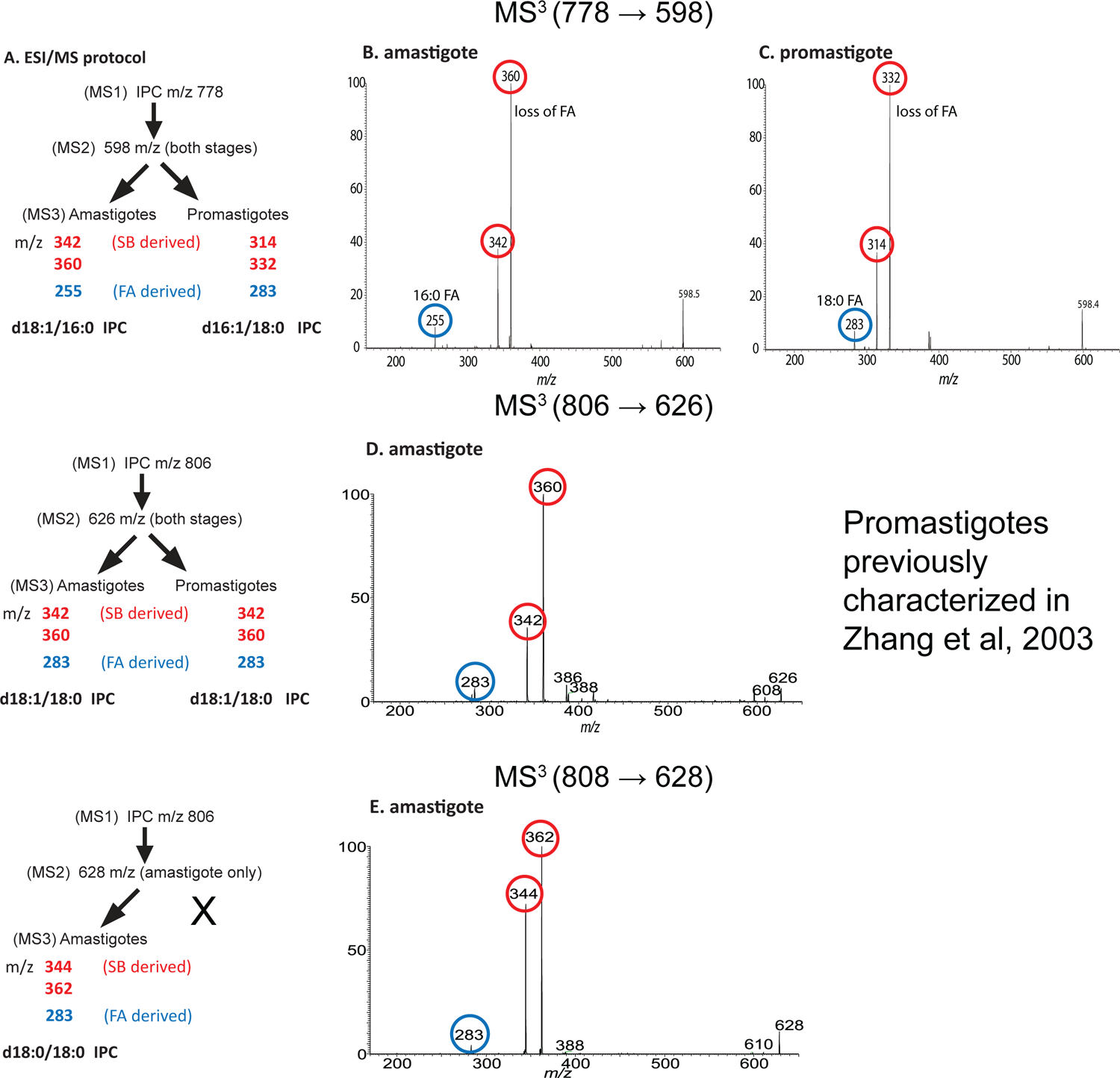
Characterization of IPC ceramide anchors. (A), Summary of ESI/MS fragmentation results and deduced ceramide anchor structures. (B,C), MS^3^ (*m/z* 778.6 → *m/z* 598) analysis of the IPC ceramide moiety of *m/z* 598 from amastigotes (B) or promastigotes (C). Unlike promastigote d16:1/18:0-IPC, the deduced amastigote IPC anchor is the same as that of mammalian d18:1/16:0-SM. (D), MS^3^ (*m/z* 806.6 → *m/z* 626) analysis of the IPC ceramide moiety of *m/z* 626 from amastigotes defined the d18:1/18:0-IPC structure, identical to that previously reported for promastigotes (32). (E), MS^3^ (*m/z* 808.6 → *m/z* 628) analysis of the amastigote IPC ceramide moiety of *m/z* 628 (not seen in promastigotes) defined the d18:0/18:0 IPC structure. Fragment ions derived from the fatty acid moiety are highlighted or circled in blue, while SB-derived moieties (acid or ketene as described in the text) are highlighted or circled in red.

For the amastigote IPC *m/z* 806.6 species, the MS^2^ spectrum contained the ion *m/z* 626 arising from loss of inositol (Fig. 9A), which was further dissociated (MS^3^) to ions of *m/z* 360 and 342 by loss of 18:0-FA as ketene and acid, respectively. The spectrum also contained the ion of *m/z* 283, representing a 18:0 fatty acid carboxylate anion (Fig. 9D). The combined structural information led to define the d18:1/18:0-IPC structure, which is identical to that previously determined for the promastigote IPC (*m/z* 806.6) (32). Lastly, MS^2^ on the unique amastigote IPC ion of *m/z* 808.6 gave rise to an ion of *m/z* 628 (loss of inositol) (Fig. 9A), which was further dissociated (MS^3^) to ions of *m/z* 362 and 344 by loss of 18:0-FA as ketene and acid respectively, along with the 18:0 carboxylate anion of *m/z* 283 (Fig. 9E). These results readily defined a d18:0/18:0-IPC structure.

The above structural information of IPCs from amastigote and promastigote (summarized in Fig. 9A) indicates that IPC synthesis in amastigotes may arise following salvage and further metabolism, with two unique ceramide anchors not seen in promastigote IPCs. and is considered further in the discussion.

## Discussion

Here we characterized IPC synthesis by *Leishmania major* across the infectious cycle, and through genetic deletion established a surprising lack of impact of IPC and *IPCS* knockouts on parasite biology and virulence in susceptible mouse infections. Our data suggest this likely reflects extensive salvage of host sphingolipids including sphingomyelin. We consider these findings in the context of SL uptake and metabolism, and their consequences towards efforts to target parasite SL metabolism in chemotherapy.

### Minimal impact of *IPCS* deletion on promastigotes *in vitro*

In contrast to other eukaryotic microbes, *IPCS*-null (*Δipcs*^-^) parasites could be readily obtained (Fig. 1), which as promastigotes appeared nearly normal, with only small differences in cell shape, growth and viability (Fig. 4A,B). *Δipcs*^-^ promastigotes lacked IPCS enzymatic activity (Figs. 2, S1), and both promastigotes and amastigotes lacked IPC (Figs. 3, 6), ruling out alternatives sources of IPC synthesis across the infectious cycle. We also confirmed that endogenously synthesized IPCS was insensitive to aureobasidin A (Fig. 2), as seen with recombinant enzyme and mitigating a role for hypothetical IPCS-interacting proteins (26, 49). In all studies, ectopic re-expression of *IPCS* restored IPC synthesis and all other phenotypes tested (Figs. 3-5). IPCS activity consumes ceramide and PI to yield IPC and DAG, however the health of *Δipcs*^-^ suggests that perturbations of these substrates in this context likely has little deleterious effect during the infectious cycle. Provision of high levels of exogenous DAG had little effect on *Δipcs*^-^ but feeding SBs proved toxic, presumably due to excessive accumulation of SBs and/or ceramide in the absence of further metabolism, as seen in *Leishmania* and fungi (38, 50). *Δipcs*^-^ was strongly inhibited relative to WT when grown at pH 5 (Fig. 4D), however this effect was largely ameliorated at pH 5.5 or above (Supplemental Table 2). This is consistent with the survival of *Δipcs*^-^ in animal infections (Fig. 5), as the *Leishmania major* parasitophorous vacuole has a pH of 5.4 (51) which would only weakly impact *Δipcs*^-^. In these properties, *Leishmania* differs significantly from *Cryptococcus neoformans*, where *IPCS* is essential and IPCS-deficient organisms are hyper-susceptible to pH stress through an absence of DAG signaling (22).

### Minimal impact of *IPCS* deletion on virulence and pathology

Distinct from other eukaryotic microbes, *Δipcs*^-^ *L. major* showed no defects in virulence as revealed by their ability to survive and induce progressive pathology when inoculated into susceptible mice (Figs. 5, S4A), consistent with the findings *in vitro* with promastigotes where there was little change in properties expected to play significant role in amastigotes. In fact, lesion pathology was somewhat enhanced in the *Δipcs*^-^, as was the parasite burden (Figs. 5, S5).

Given the relative health and virulence of amastigotes lacking IPC, the question of why *<u>Leishmania</u>* continue to synthesize it at all now arises. Perhaps rather than contributing to bulk SL levels or metabolism, specific recognition of IPC by host defenses could contribute to *Leishmania* pathology in some circumstances not tested here. Several studies have reported interactions between *Leishmania* and host ceramide impacting macrophage survival of the related species *L. donovani* (52, 53). We did observe somewhat higher levels of ceramides in purified amastigotes relative to infected host tissue (Fig. S6), and speculatively the modest hypervirulence seen with *L. major Δipcs*^-^ might arise from this.

### Abundant salvage leads to predominance of SM over IPC in WT *Leishmania* amastigotes: a potential role in virulence

*Leishmania* promastigotes can survive perfectly well without any cellular SLs (31, 32), raising the possibility that the amastigote stage could likewise lack a requirement for SLs. However, *Leishmania* sp. are known to take up a variety of host lipids (54–56), and perhaps potential amastigote-stage SL requirements could be satisfied through uptake and metabolism of host SLs. MS analysis of purified amastigotes showed high levels of host SM in both WT and *Δipcs***^-^**, with SM estimated to be about 7-fold more abundant than parasite IPC (estimated previously to be present at about 10^8^ molecules/amastigote) (34). Thus host SM represents by far the most abundant salvaged SL within the parasite, even in WT amastigotes where it greatly predominates over IPC. This ‘redundancy’ suggests that loss of IPC in *Δipcs*^-^ might confer only a slight reduction in total SLs, which might have little effect or be ameliorated by a small increase in SM salvage.

That amastigotes salvage high levels of host SLs could be taken as evidence that these are required for parasite survival and virulence in the mammalian host, especially given that most eukaryotes require SL. However, given promastigotes ability to survive without SLs, potentially amastigotes could do likewise. To test this definitively, some approach where SL acquisition is disrupted chemically or by genetic mutation would be most informative, but at present none of these tools are available in *Leishmania*. We favor the hypothesis that like lipid salvage generally, SL acquisition is likely to play a key role in virulence (54,57,58).

### Metabolism of salvaged host SLs by *Leishmania* amastigotes

While the majority of acquired SM remains intact in amastigotes, that further metabolism of acquired host SLs leading to parasite IPC synthesis occurs is mandated by the fact that *de novo* sphingoid base synthesis is shut down in amastigotes, a finding recapitulated in Δ*spt2^-^* amastigotes lacking serine palmitoyl transferase (32, 34). Not only are IPC levels maintained in amastigotes, but amastigotes bear two IPCs with ceramide anchors not seen in promastigotes (d18:1/16:0-IPC and d18:0/18:0-IPC), with alterations in relative abundance as well (Fig. 9).

Several studies of murine host macrophages show that they elaborate a spectrum of SLs bearing anchors that could serve as a source for those present in amastigote IPC (47, 48) (Table S3; a summary of these data is presented in Fig. 10). It can be seen that potential donors for the three amastigote IPC ceramide anchors can be found in both host SM and ceramides, or even through various SBs and ceramide synthesis (Fig. 10; the shading is based on the amastigote IPC anchor structures). Other SL possibilities may include host glucosylceramides which were not considered further here.

**Figure 10:**
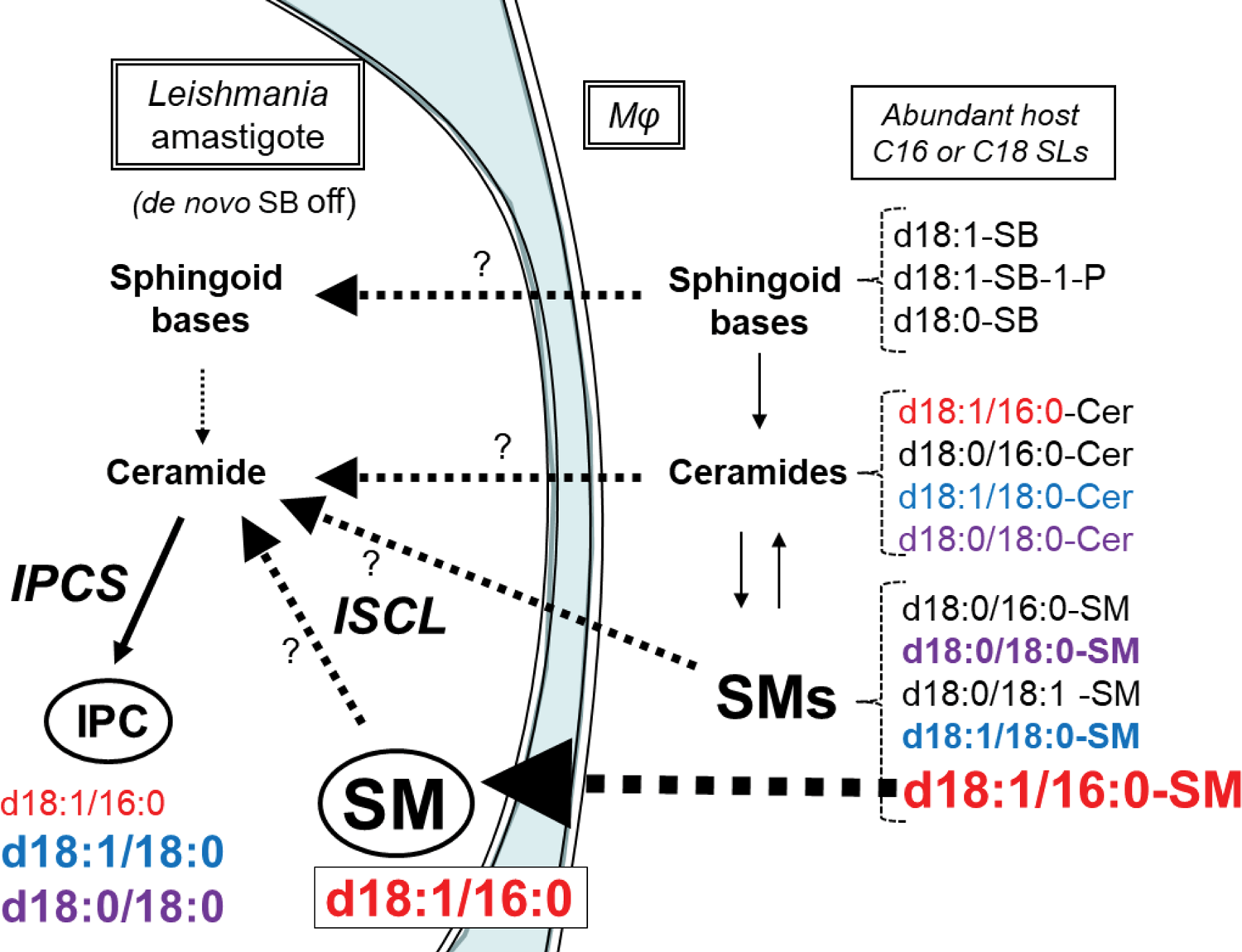
Summary of relevant SLs and their relative abundances in *Leishmania* amastigotes and murine host macrophages, and potential salvage and metabolic routes. *Leishmania* amastigotes lack *de novo* sphingoid base (SB) synthesis so SL synthetic and/or salvage pathways are shown starting at this point. The spectrum of SLs considered is restricted to SLs with C16 and C18 fatty acyl chains relevant to IPC synthesis, and is based on the present study or published data (47, 48) (Table S3). The structures of ceramides or the ceramide anchor of SMs and IPCs is shown, with larger font heights signifying approximate relative abundance. The host ceramide or SM anchors are colored to match those of the *Leishmania* IPCs to facilitate discussion about potential host origins. Question marks indicate lack of information about the relative role of the indicated pathway in IPC synthesis. The heavy dashed black arrow marks the highly abundant d18:1/16:0-SM in both *Leishmania* amastigotes and the host, and is discussed further in the text.

Focusing first on SM donors because of the massive uptake of host d18:1/16:0-SM, one potential route is through the potent sphingomyelinase / inositolphosphorylceramidase activity encoded by *Leishmania ISCL* (45,57,59,60) (Fig 10). Alternatively (and not exclusively), an *ISCL*-independent route could involve salvage of mammalian SBs or ceramides with related anchors directly (Fig. 10). Both routes are challenged by the *a priori* expectation that the ratio of the various amastigote IPC anchors would be expected to mirror that of the salvaged SL forms, from either *ISCL*-dependent or independent routes. However, while amastigote d18:1/16:0 IPC is the least abundant IPC isoform, constituting less than 20% of total IPC, d18:1/16:0 SM is by far more abundant than other isoforms, constituting more than 80%, in either the parasite or the host (47, 48) (Fig. 7, S7; Table S3). Factors that could contribute to these differences include differences in relative uptake of different SMs or ceramides, preferential salvage of host ceramides rather than ISCL-mediated degradation of SMs, or intrinsic differences in these activities with SMs or ceramides of differing anchor structure, for which there is currently little knowledge. Alternatively, an *ad hoc* hypothesis postulates there are two pools/routes of SL uptake, one emphasizing the abundant d18:1/16:0 SM, and the other more broadly acting across different IPC or ceramide anchor classes feeding into IPC synthesis (Fig. 10). Future studies will be required to address these interesting questions.

Interestingly and unlike *de novo* SL or IPC synthesis, the *ISCL* SMase activity is localized to the parasite mitochondrion where it is required for *Leishmania* virulence, albeit with some variation amongst parasite species or hosts (45, 59). Beyond bulk catabolism, the fungal orthologs of *ISCL* play roles in mitochondrial metabolism, replication and SL signaling (61, 62). Currently the molecular mechanism(s) by which SM or other SLs are acquired or partitioned into various metabolic pathways by *Leishmania* amastigotes is unknown, and unfortunately genetic approaches have been uninformative since no *Leishmania* mutants lacking SLs entirely have emerged.

### Implications of the relative ‘insignificance’ of IPCS to chemotherapy

We did not anticipate that IPC or IPCS would not be essential at *any* point in the *Leishmania* infectious cycle, unlike pathogenic fungi where essential IPCS shows considerable potential as a validated, drug target. In this respect, our data do not provide much optimism for IPCS as a drug target in the *Leishmania* parasite. While the yeast IPCS inhibitor aureobasidin A inhibits *Leishmania* growth or infectivity at relatively high concentrations (>10 µM) (25, 63), this was considered ‘off-target’ as the susceptibility was unchanged when tested against *L.major Δspt2***^-^** mutants completely lacking SLs (25). Off-target effects were also observed in the related kinetoplastids parasite *Trypanosoma cruzi* and the apicomplexan parasite *Toxoplasma gondii* (64, 65). Most importantly, all data show that aureobasidin A is inactive against recombinant IPCS from trypanosomes and *Leishmania,* (25, 26) as well as microsomal *Leishmania* IPCS activity (this work, Fig. 2). Whether new candidate inhibitors showing activity against cultured parasites (26–29) act ‘on target’ against *Leishmania* IPCS remains to be seen, although the health of the *Δipcs*^-^ mutant suggest this may be unlikely. *In vitro*, IPCS inhibitors may act synergistically with other compounds, although whether this extends to the *in vivo* situation with abundant SL salvage remains to be seen, in *Leishmania* or other pathogens known to take up host lipids.

Our studies thus provide a new perspective on *Leishmania* SL pathways, emphasizing the role of SM salvage from the host cell over *de novo* synthesis by the amastigote stage, and additionally raising many interesting questions that will require further study. Instead, our findings suggest that agents inhibiting SL uptake or catabolism in *Leishmania* may have more promise as an avenue for chemotherapy.

### Experimental Procedures

#### Molecular constructs

Molecular constructs containing 610 bp of 5’ flanking (PCR amplified from WT genomic DNA by primers SMB 2590 and 2591; Table S1 contain the sequences of all oligonucleotides used) and 665 bp of 3’ flanking (amplified by primers SMB 2592 and 2593) sequence flanking the *IPCS* ORF were joined by fusion PCR to the ORF for the selectable markers NEO (amplified by primers SMB2586 and 2587) or SAT (amplified by primers SMB 2588 and 2589). These constructs (pGEM T Easy-IPCS::ΔNEO, B5810, or pGEM T Easy-IPCS::ΔSAT, B6075) were linearized by digestion with EcoRI to expose the homologous ends prior to electroporation. To restore IPCS expression, the IPCS coding region (Lm35.4990) was amplified by PCR using the primers SMB 3259 and 3260 and inserted into the pXG expression vector (66, 67) yielding pXG-BSD-IPCS (B6235). All constructs were confirmed by sequencing.

#### *Leishmania* culture and transfection

*Leishmania major* (LV39 clone 5, Rho/SU/59/P) was grown in M199 media supplemented with 10% fetal bovine serum, 12 mM NaHCO_3_, 0.1 mM adenine, 7.6 mM hemin, 2 μg/ml biopterin, 20 μM Folic Acid, 1x RPMI Vitamin Mix, 2mM glutamine, 25 mM HEPES pH 6.8, 50 U/ml penicillin, and 50 μg/ml streptomycin (68). Cell density was measured using a coulter counter (Becton Dickinson) and viability was determined by flow cytometry (Becton Dickinson) after staining with 5μg/ml propidium iodide. In some experiments supplements were added as indicated, or the pH was altered as shown in table S1 and MES or citrate buffers. Metacyclic cells were prepared by density purification or PNA-purification as described (40).

Transfections were performed by electroporation as previously described (69). To replace the first IPCS allele, transfection with the *IPCS:: ΔNEO* targeting fragment was performed; due to some media/serum concerns at the time these studies were undertaken, we had difficulty recovering transfectant colonies by our usual plating procedures. Thus, after electroporation and a 24 hr recovery period in 3 separate pools, clonal lines were obtained by limiting dilution in M199 medium containing 10 μg/ml neomycin and 500 μM EtN. The authenticity of several putative heterozygous transfectants (*IPCS / IPCS:: ΔNEO*) was confirmed by PCR. Correct integration was confirmed by using primers outside the flanking region and inside the drug marker (SMB2598/2600 5’Flank and SMB2608/2827 3’Flank). Several independent clonal lines were inoculated into susceptible BALB/c mice (10^7^ parasites/footpad) and recovered after 1 month; such lines are referred to as M1 passage. Two such lines (WA and WB) were then transfected with the *IPCS:: ΔSAT* targeting fragment, resistant cells obtained following growth in 8 μg/ml neomycin and 100 μg/ml nourseothricin, and clonal lines obtained by limiting dilution. The authenticity of potential homozygous Δ*ipcs^-^* null mutants (formally *IPCS:: ΔNEO/IPCS:: ΔSAT*) was confirmed by PCR as described above using primers SMB 2598/2601 (5’ flank) and SMB 2609/2827 (3’ Flank). Several lines were inoculated into BALB/c mice as before. IPCS expression was restored in several Δ*ipcs^-^* M1 transfectants (WAB, WAD, and WAE) by transfection with pXG-BSD-IPCS (B6235) to generate *ipcs::ΔNEO/ipcs:: ΔSAT/+pXG-IPCS*, referred to as Δ*ipcs^-^*/+*IPCS*. As all transfectant and add-back lines showed similar behavior in preliminary tests, detailed studies are presented only for clone WAD (Δ*ipcs^-^*) or WADB (Δ*ipcs^-^/+IPCS*) unless otherwise stated.

### Microsomal Preparation and Enzyme Assays

Microsomes were prepared by washing 4 x 10^8^ promastigotes in PBS and homogenized by sonication in buffer (50mM MES, 50 mM BIS-TRIS pH 6.25, 10 mM EDTA, 100 mM NaCl, 250 mM Sucrose, 300 nM aprotinin, 1 mM benzamidine, 1μg/ml chymostatin and pepstatin, 10 μM E64 and leupeptin, 10 mM NaF). Debris was removed by centrifugation at 1,000 x g for 10 min and the supernatant was then centrifuged at 100,000 x g for 90 minutes. The pellet was resuspended in minimal amounts of buffer (5 mM MES, 5 mM BIS-TRIS pH 6.25, 1 mM EDTA, 100 mM NaCl, 15 % glycerol, 300nM aprotinin, 1mM benzamidine, 1μg/ml chymostatin and pepstatin, 10 μM E64 and leupeptin, 1 mM NaF) with sonication then flash frozen. Protein content was determined by BCA assay using BSA standards.

Enzyme assays were performed in 100 μl IPCS assay buffer (100 mM MES, 100 mM BIS-TRIS pH 6.25, 5mM EDTA, 100 mM NaCl, 300 nM aprotinin, 1 mM benzamidine, 1 μg/ml chymostatin and pepstatin, 10 μM E64 and leupeptin, 1 mM NaF). Lipid substrates were dried by speedvac prior to resuspension is assay buffer, using 625 picomoles of L-α-phosphatidylinositol from soy and 250 picomoles of NBD-C_6_-Ceramide unless otherwise indicated. 25 μg of microsomal protein was added to the reactions which were incubated at 28°C for 60 minutes and terminated with 500 μl of 50 mM HCl in methanol. Where indicated aureobasidin A was added to a final concentration of 20 µM. Lipids were extracted with 1.5 ml of chloroform and 463 μl of 1M NaCl. Samples were vortexed and centrifuged at 1,500 x g for 5 min. The upper phase was re-extracted with 750 μl of chloroform:methanol (43:7) and the combined lower phases were dried by nitrogen stream. Samples were resuspended in 50 μl of chloroform:methanol (2:1) and 5 μl were loaded on predeveloped Baker Si-250-PA 60 A silica gel TLC plates and developed using chloroform:methanol:water:ammonium hydroxide (28%)/65:30:3:2/v:v:v:v. Images were developed using Fuji FLA-5000 phosphoimager.

### Lipid Extraction

Promastigotes were collected from 100 ml cultures and lesion amastigotes were harvested as described (70). Sphingolipids from log phase promastigotes or amastigotes were extracted using a modified Folch extraction. First, 2 x 10^8^ cells were resuspended in 1.5 ml 25 mM Sodium Fluoride with 2 ml Methanol and sonicated. Next, 4 ml of chloroform was added; samples were vortexed and centrifuged at 1,800 x g for 15 minutes. The upper phase was re-extracted with 2.5 ml of chloroform:methanol (43:7), vortexed, and centrifuged. Combined lower phases were back extracted with 2 ml of 50 mM sodium fluoride and 2 ml of Methanol, vortexed, and centrifuged. The lower phase was dried by nitrogen stream and extracted again with chloroform:methanol:50 mM sodium fluoride (4:2:1), vortexed, and centrifuged. The lower phase was again dried by nitrogen stream and resuspended in 1 ml of chloroform:methanol (1:1) for analysis by mass spectrometry.

### Mass Spectrometry

ESI/MS analyses were performed on a Thermo Scientific (San Jose, CA) triple stage quadrupole (TSQ) Vantage EMR mass spectrometer with Xcalibur operating system. Lipid extracts (10 uL each injection) were injected to the HESI-II probe in a Thermo Scientific™ Ion Max ion source by a built-in syringe pump which delivered a constant flow of methanol at 20uL/min. The mass spectrometer was tuned to unit-mass resolution, and the heated capillary temperature was set to 280°C. The electrospray needle was set at 4.0 kV and 3.5 kV for positive-ion and negative-ion modes operation, respectively, and the skimmer was at ground potential. The mass resolution of both Q1 and Q3 was set at 0.6 Da at half peak height to obtain the product-ion scan, precursor ion scan and neutral loss scan (NLS) mass spectra, using a collision energy of 30-40 eV with target gas of Ar (1.3 mtorr) in the rf-only second quadrupole (Q2). The mass spectra were accumulated in the profile mode. Significant peaks were assigned based on signal to noise ratio to assess significance or quantitation (43), and evaluated for natural isotope patterns (44).

### Mouse Virulence

Mouse handling and experimental procedures were carried out in strict accordance with the recommendations in the Guide for the Care and Use of Laboratory Animals of the United States National Institutes of Health. Animal studies were approved by the Animal Studies Committee at Washington University (protocol #20-0396) in accordance with the Office of Laboratory Animal Welfare’s guidelines and the Association for Assessment and Accreditation of Laboratory Animal Care International.

Briefly, BALB/c or γ-IFN knockout C57BL/6 mice (γKO) were infected with 10^5^ metacyclics purified by gradient centrifugation (40), and footpad lesion swelling measured weekly. At key points parasites were enumerated by limiting dilution assays (71), or lesions were excised for purification of amastigotes as described previously (34).

### Statistical Analysis

Statistical analysis was performed indicated using GraphPad Prism version 9.4.1

## Supporting information

This article contains supporting information available on-line.

## Acknowledgments

We thank the members of our laboratory and Kai Zhang (Texas Tech University, Lubbock) for helpful discussions and providing access to MS data of purified WT amastigotes for confirmation of the results presented here. This manuscript was submitted posthumously on behalf of John Turk.

## Author contributions

FMK, PK, FH and SMB, conceptualization; FMK, PK, SH and FH, investigations, FMK, FH and SMB, writing; and JT and SMB, resources, supervision and funding.

## Abbreviations

WT: wild-type *Leishmania major*

SM: sphingomyelin

IPC: inositolphorphorylceramide

IPCS: inositolphosphorylceramide synthase

SL: sphingolipid

SLS: sphingolipid synthase

SB: sphingoid (long-chain) base

PC: phosphatidyl choline

PI: phosphatidyl insotiol

PI-PLC: phosphatidyl inositol phospholipase C

NBD-Cer: (6-((N-(7-Nitrobenz-2-Oxa-1,3-Diazol-4-yl)amino)hexanoyl)Sphingosine)

MS: mass spectrometry

ISCL: inositol phosphorylceramide phospholipase C-like

ESI/MS: electrospray ionization mass spectrometry.

γKO: γ-interferon knockout C57BL/6 mice

SD: standard deviation.

Lipid species abbreviations follow those recommended by the Lipid Maps consortium (www.lipidmaps.org).

## Supporting information

### Materials included

**Figure S1:**
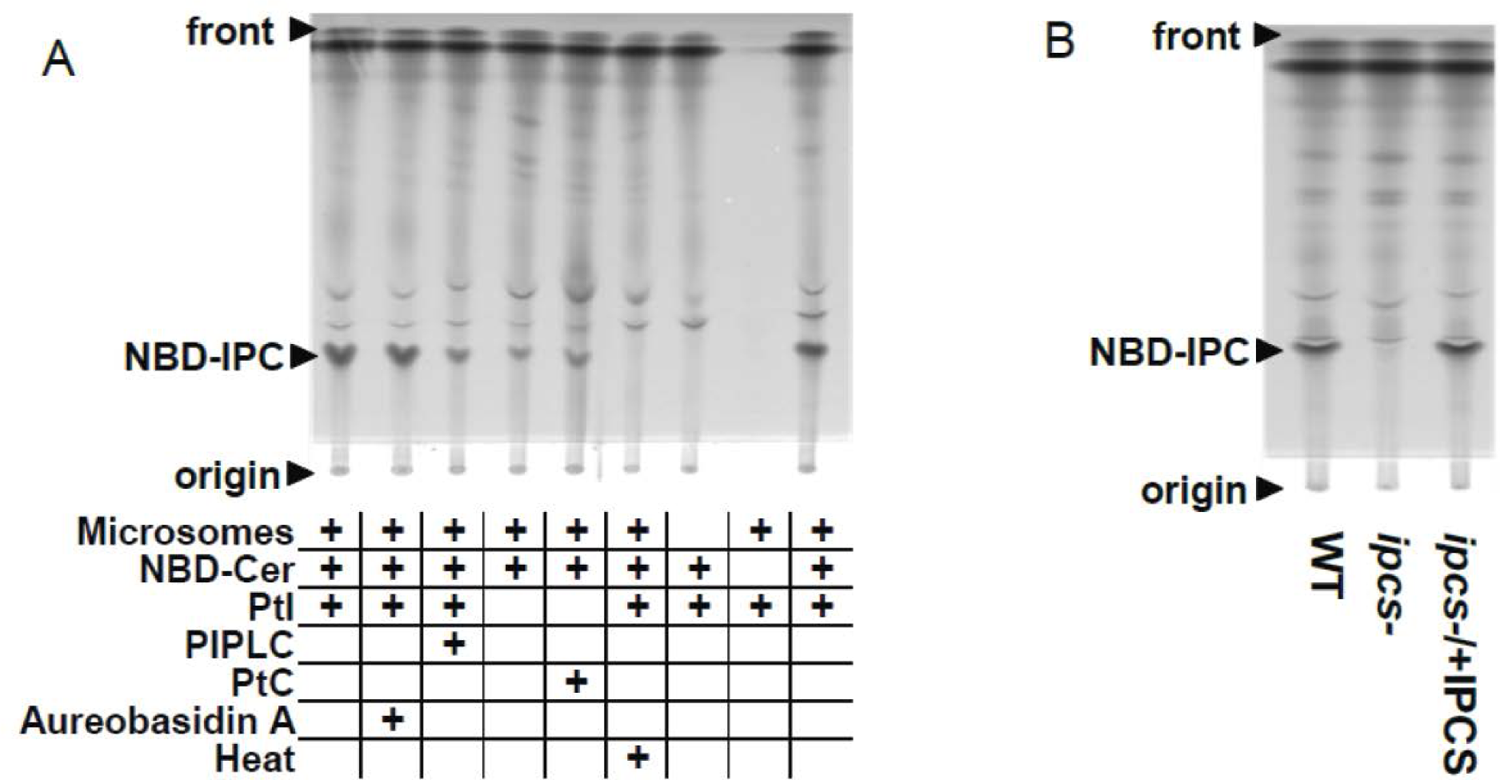
Enzymatic characterization of IPCS activity from purified microsomes, and its absence in Δ*ipcs^-^* promastigotes. The figure displays representative TLC separations used for the quantitative plots shown in Fig. 2.

**Figure S2:**
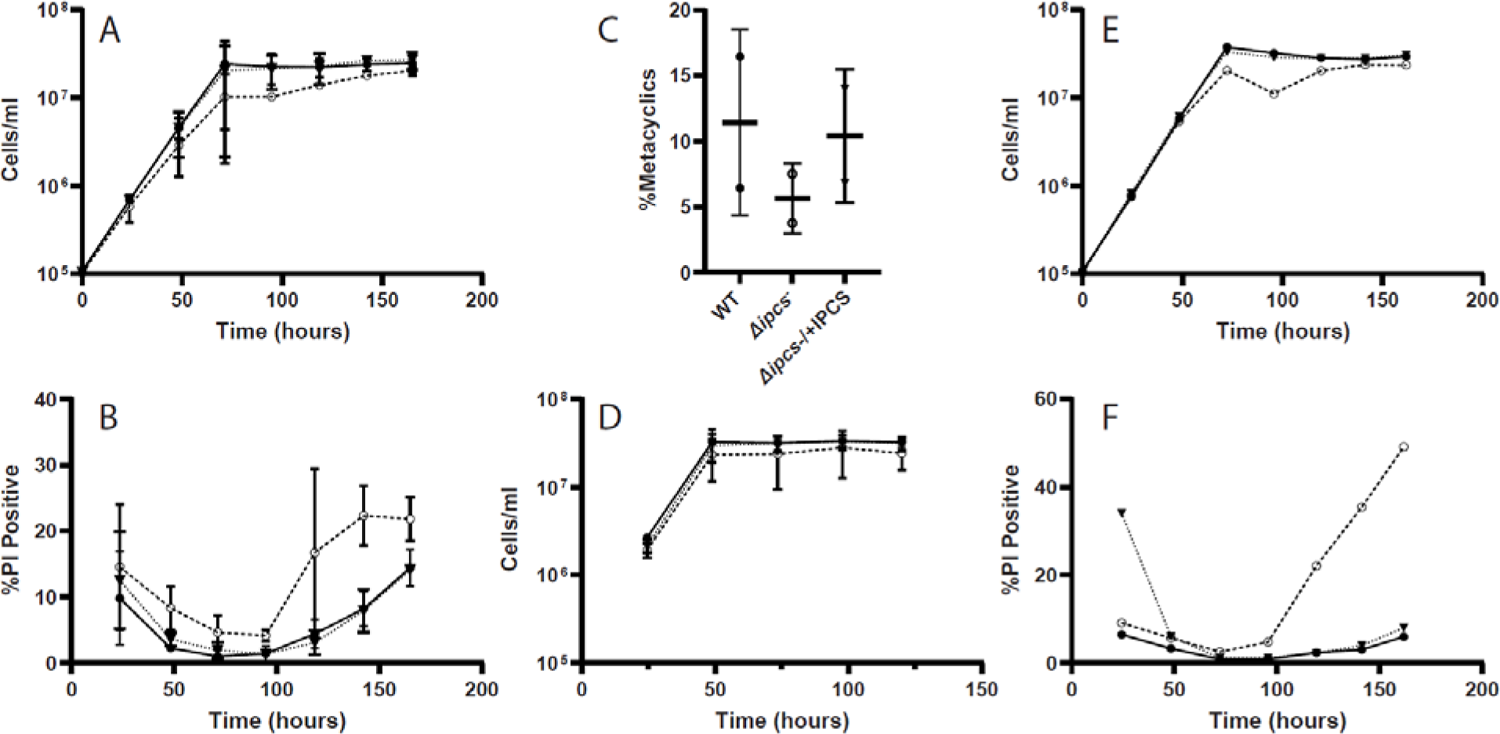
Phenotypic characterization of WT and Δ*ipcs^-^* mutant *L. major* propagated in M199 medium *in vitro*. (A, B), Growth and viability of parasites grown in 500 µM EtN. (C), Percent metacyclic observed at day 2 in stationary phase; (D), Growth of parasites at 33°C (at 37° all lines died). No significant differences amongst lines were seen in panels A-D (n = 2, means ± 1SD are shown; ANOVA followed by Bonferroni correction to adjust for multiple comparisons). (E, F), growth and viability of parasites supplemented daily with 2 μM DAG (n =1). Symbols are WT (●, solid line), Δ*ipcs^-^* (○), or Δ*ipcs^-^*/+*IPCS* (▾, dashed line).

**Figure S3:**
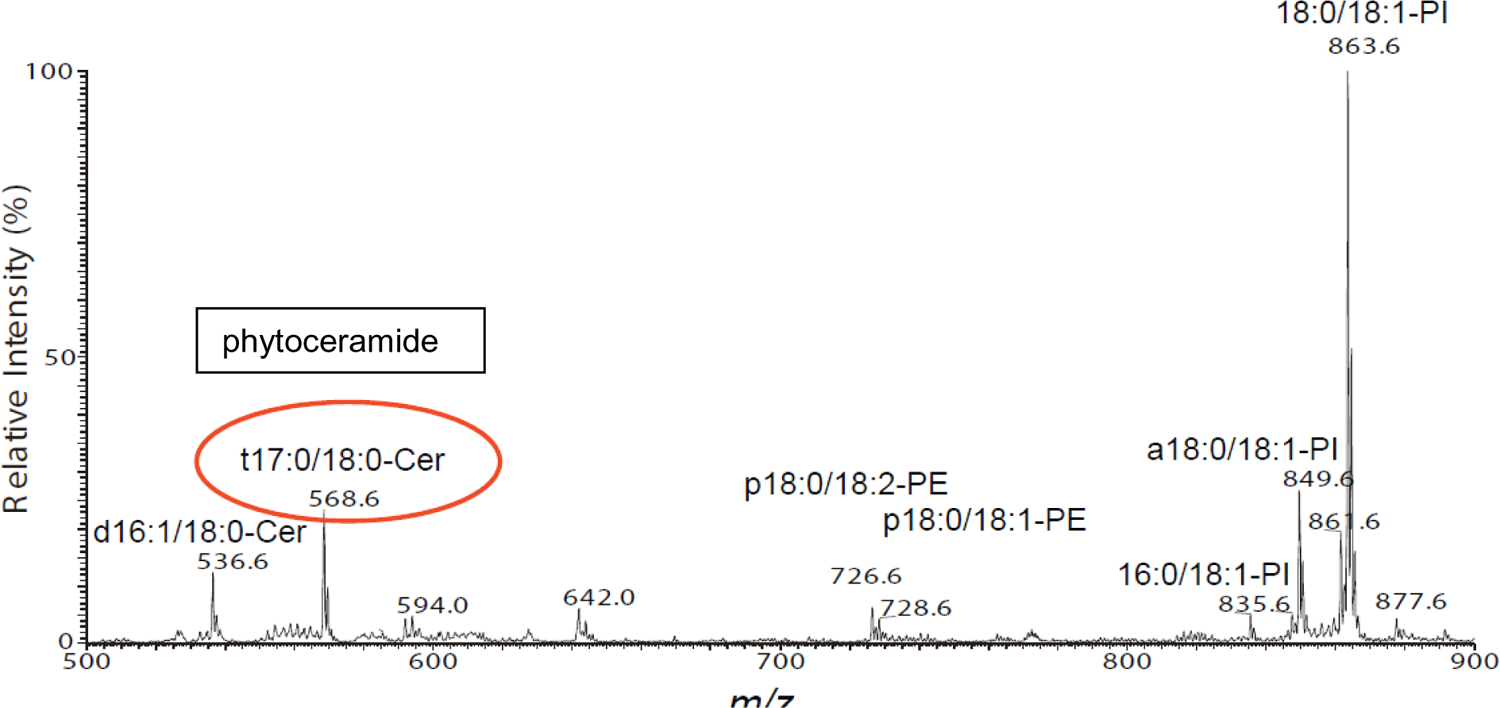
Accumulation of phytoceramide in Δ*ipcs^-^* parasites grown in the presence of phytosphingosine. The ESI/MS (negative ion mode) spectra obtained using purified lipids is shown. *In vitro* cultures of Δ*ipcs^-^* parasites were inoculated at 1 x 10^6^ cells/ml in M199 media containing 8 µM C17-phytosphingosine and grown 16 hr. The peak corresponding to phytoceramide (m/z 568) is circled. Nomenclature for individual lipid species is provided in the legends of Fig. 3 and Fig. 6.

**Figure S4:**
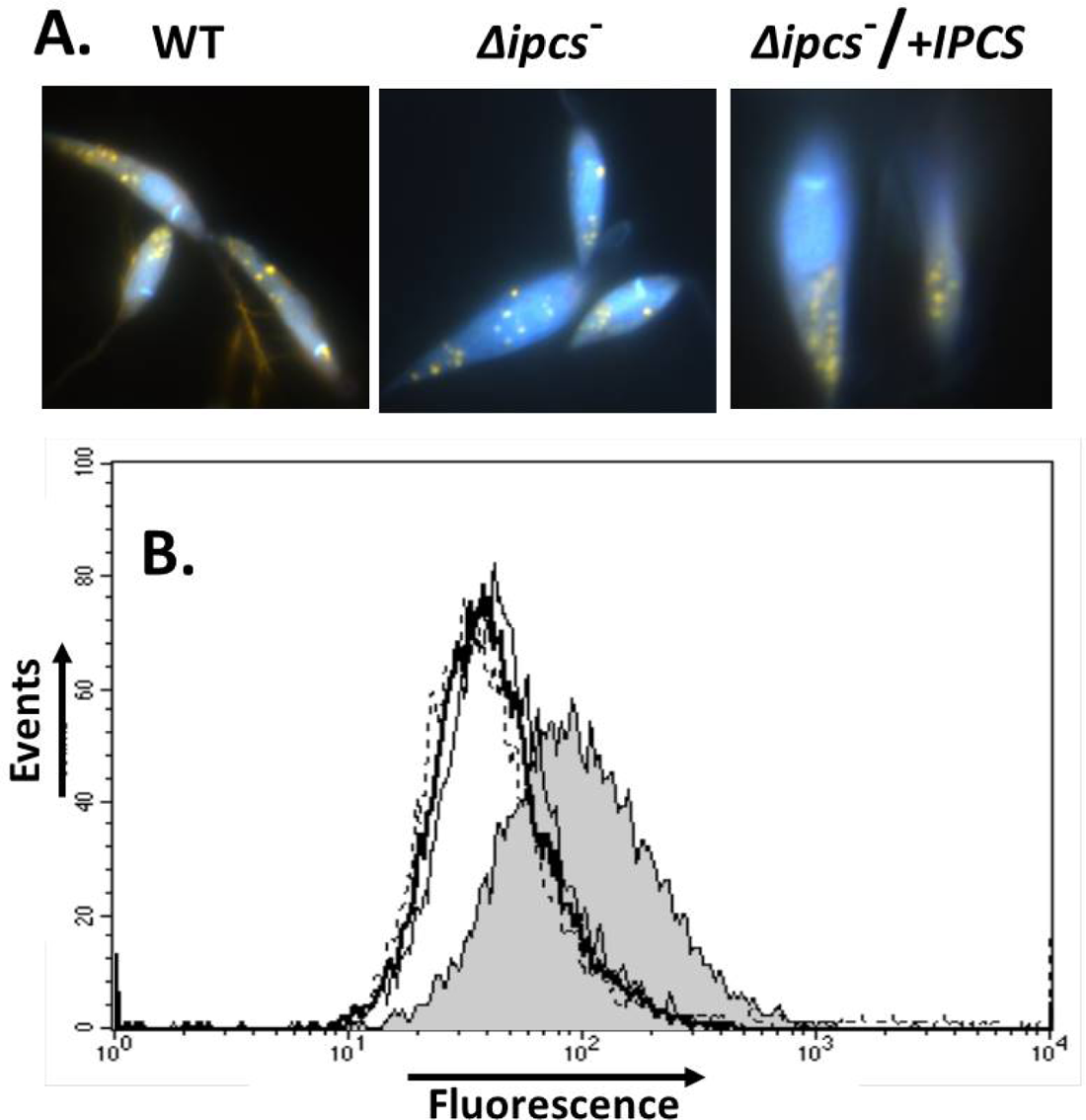
DAPI and Nile Red O staining of WT and Δ*ipcs^-^L. major*. Panel A: DAPI staining and immunofluorescence to reveal acidocalcisomes and polyphosphates (yellow) of WT, Δ*ipcs^-^*, or Δ*ipcs^-^/+IPCS*. Panel B: Flow cytometry of day stationary cells stained with Nile Red O. WT (thick solid line), Δ*ipcs^-^* (dashed line), or Δ*ipcs^-^/+IPCS* (thin solid line), all of which overlap. As a control, staining of the SL deficient mutant Δ*spt2^-^* shown previously to accumulate lipid bodies is shown (shaded area). *Methods*: DAPI staining and immunofluorescence microscopy to reveal acidocalcisomes and polyphosphates was performed as described (3, 4). Nile Red O staining for lipid accumulation (lipid bodies) and flow cytometry were performed as described (5).

**Figure S5.**
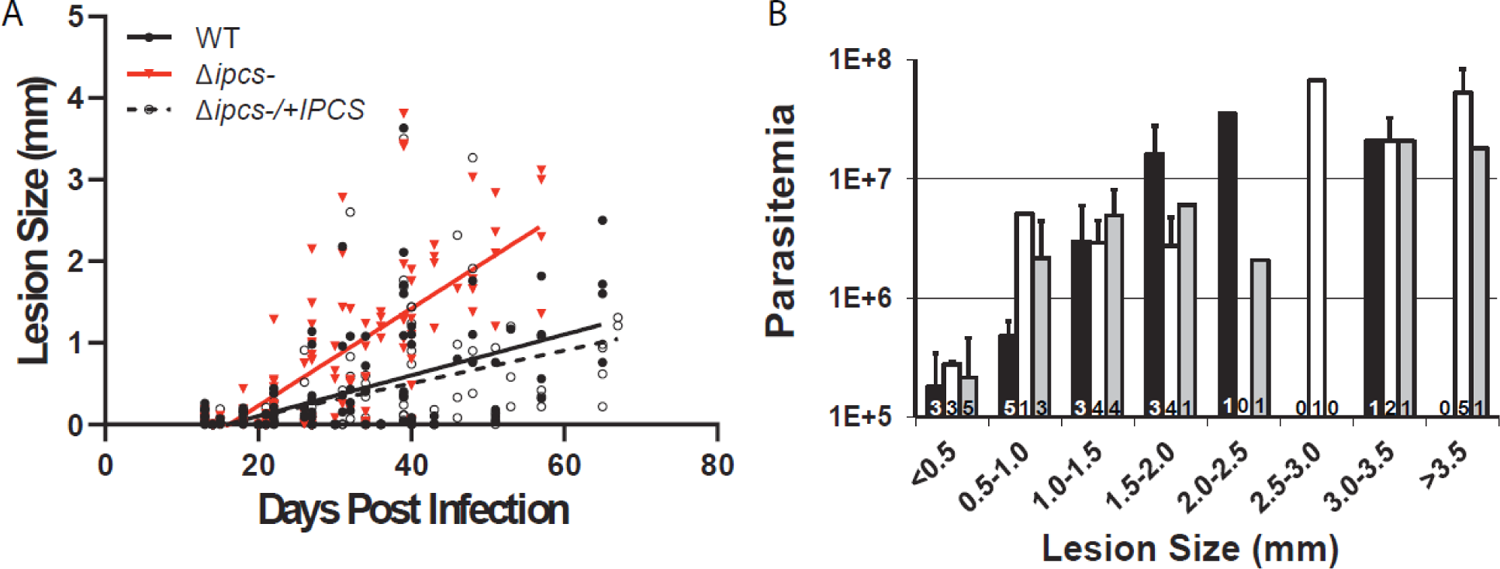
*Δipcs^-^* remains highly virulent in animal infections. (A) Collected data from four independent experiments with 4-8 mice/group are plotted (a representative individual experiment is shown in Fig. 5). The lines (slopes) shown were calculated by simple linear regression showing significant differences between *Δipcs^-^* and WT or *Δipcs^-^/+IPCS* (p < 0.0001). (B) Similar relationship between lesion size and parasite numbers in WT and mutant *L. major*. At various time points post-infection, lesions were excised and parasite numbers determined by limiting dilution. As not every experiment yielded lesions of a given size at the times measured for comparison, the small numbers within the bars denote the number of lesions, and the standard deviation is shown when possible; differences amongst lines were not considered significant. Parasite lines were WT (black bar), *Δipcs^-^* (white bar), and *Δipcs^-^/+IPCS* (gray bar) lesions.

**Figure S6.**
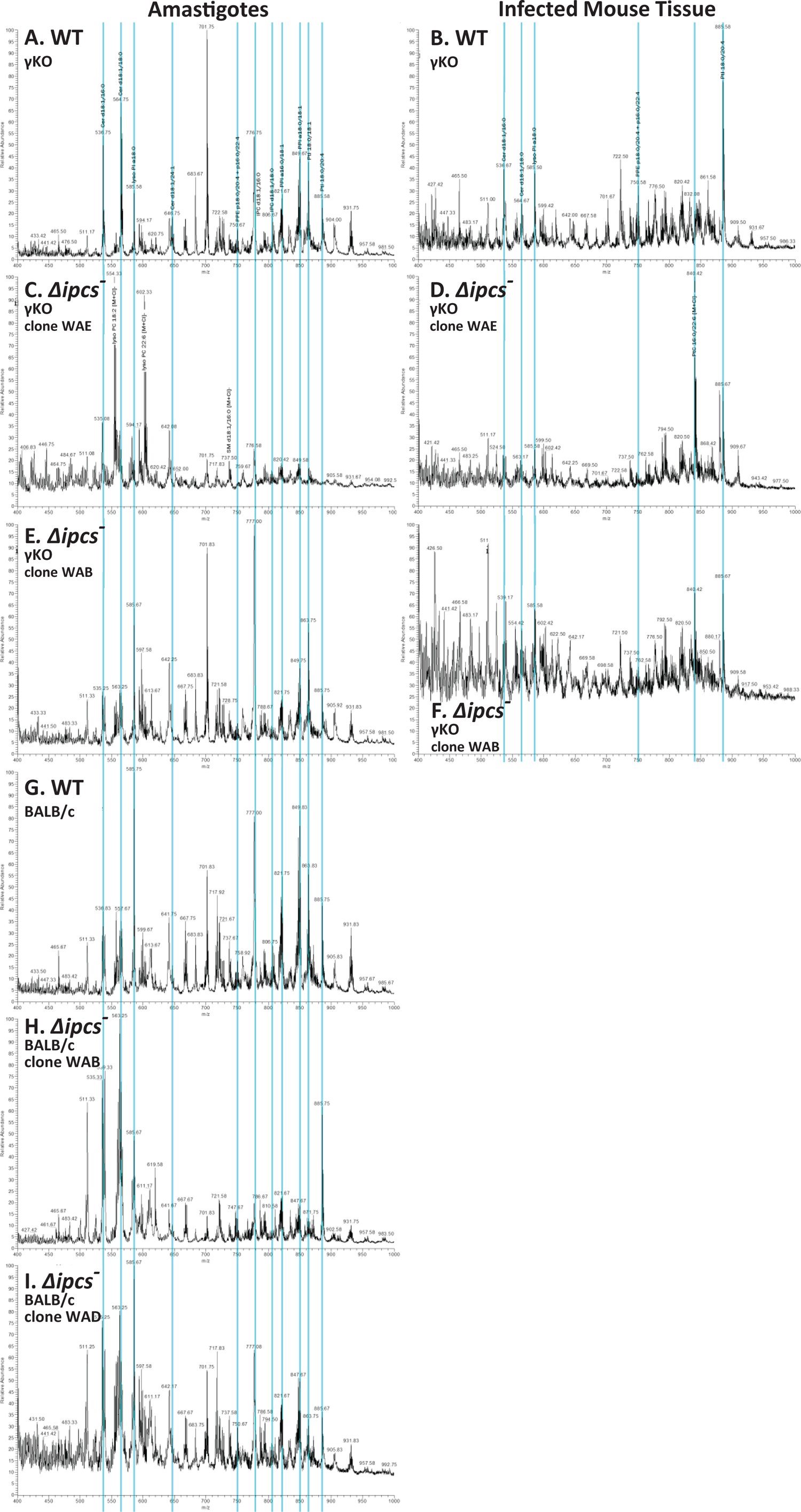
Full scan negative ion ESI mass spectra of the lipids extracted from purified amastigotes or infected mouse tissue. (A), WT amastigotes purified from γKO mice infections. (B), infected lesion tissue from WT infected γKO mice. (C), *Δipcs*^-^ amastigotes (clone WAE) purified from γKO mice infections. (D), infected lesion tissue from *Δipcs^-^* (clone WAE) infected γKO mice. (E), *Δipcs*^-^ amastigotes (clone WAB) purified from γKO mice infections. (F), infected lesion tissue from *Δipcs^-^* (clone WAB) infected γKO mice. (G), WT amastigotes purified from BALB/c mice infections. (H), *Δipcs*^-^ amastigotes (clone WAE) purified from BALB/c mice infections. (I), *Δipcs*^-^ amastigotes (clone WAB) purified from BALB/c mice infections. Major ions of interest for comparison were lined up across the panels; ions at *m/z* 808.6 (not designated) corresponds to d18:0/18.0-IPC. Lipid species abbreviations follow those recommended by the Lipid Maps consortium (www.lipidmaps.org). *Please note that due to the size, Fig. S6 can be found as a separate on-line file*.

**Figure S7.**
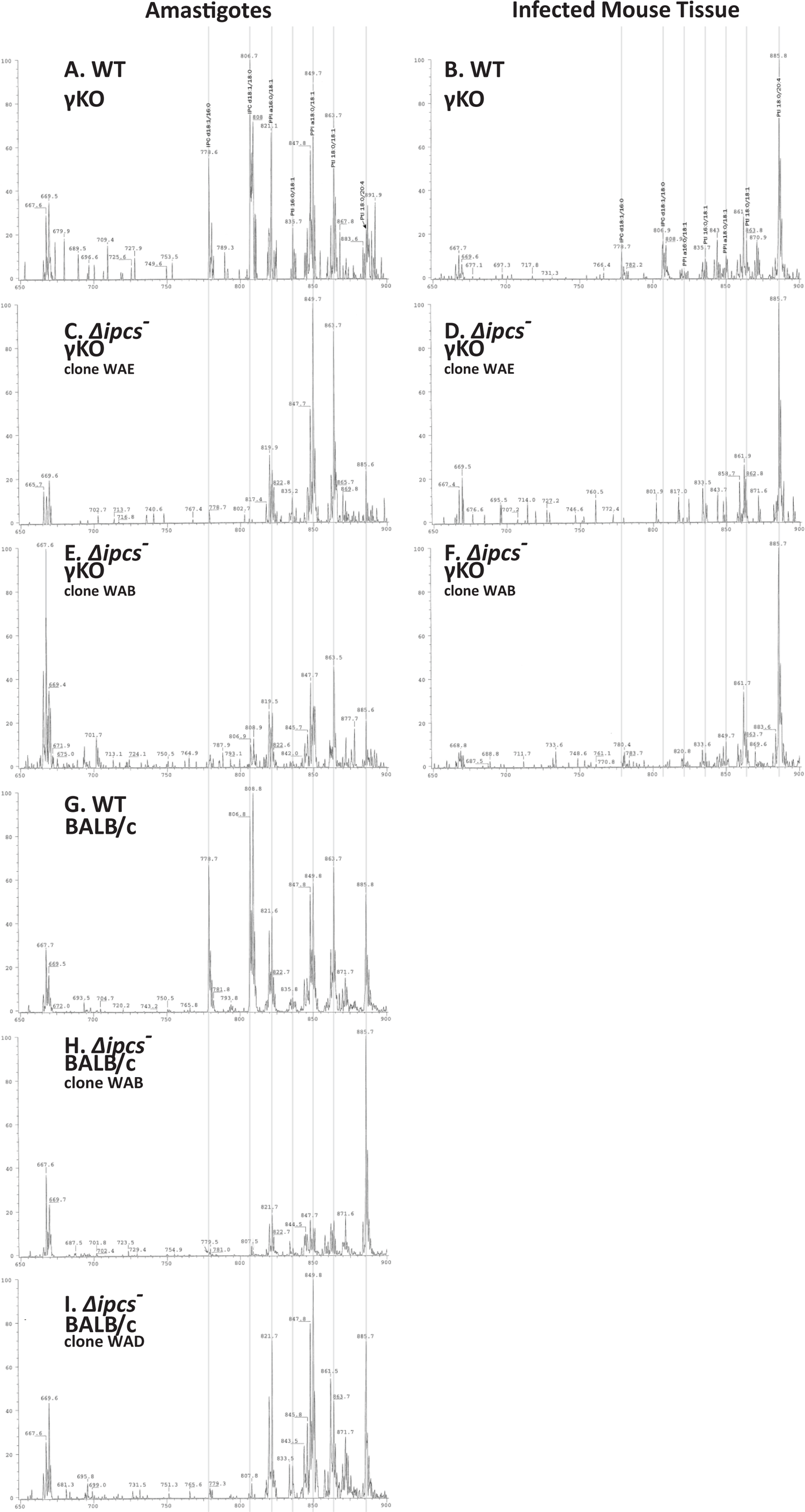
Negative ion ESI MS/MS spectra acquired by parent ion scan of 241 that detect PI and IPC in the lipid extracts from purified amastigotes or infected mouse tissue. (A), WT amastigotes purified from γKO mice infections. (B), infected lesion tissue from WT infected γKO mice. (C), *Δipcs*^-^ amastigotes (clone WAE) purified from γKO mice infections. (D), infected lesion tissue from *Δipcs^-^* (clone WAE) infected γKO mice. (E), *Δipcs*^-^ amastigotes (clone WAB) purified from γKO mice infections. (F), infected lesion tissue from *Δipcs^-^* (clone WAB) infected γKO mice. (G), WT amastigotes purified from BALB/cmice infections. (H), *Δipcs*^-^ amastigotes (clone WAE) purified from BALB/c mice infections. (I), *Δipcs*^-^ amastigotes (clone WAB) purified from BALB/c mice infections. Major PI and IPC ions were lined up across the panels for comparison; ions at *m/z* 808.6 (not designated) corresponds to d18:0/18.0-IPC. Lipid species abbreviations follow those recommended by the Lipid Maps consortium (www.lipidmaps.org). *Please note that due to the size, Fig. S7 can be found as a separate on-line file*.

**Figure S8.**
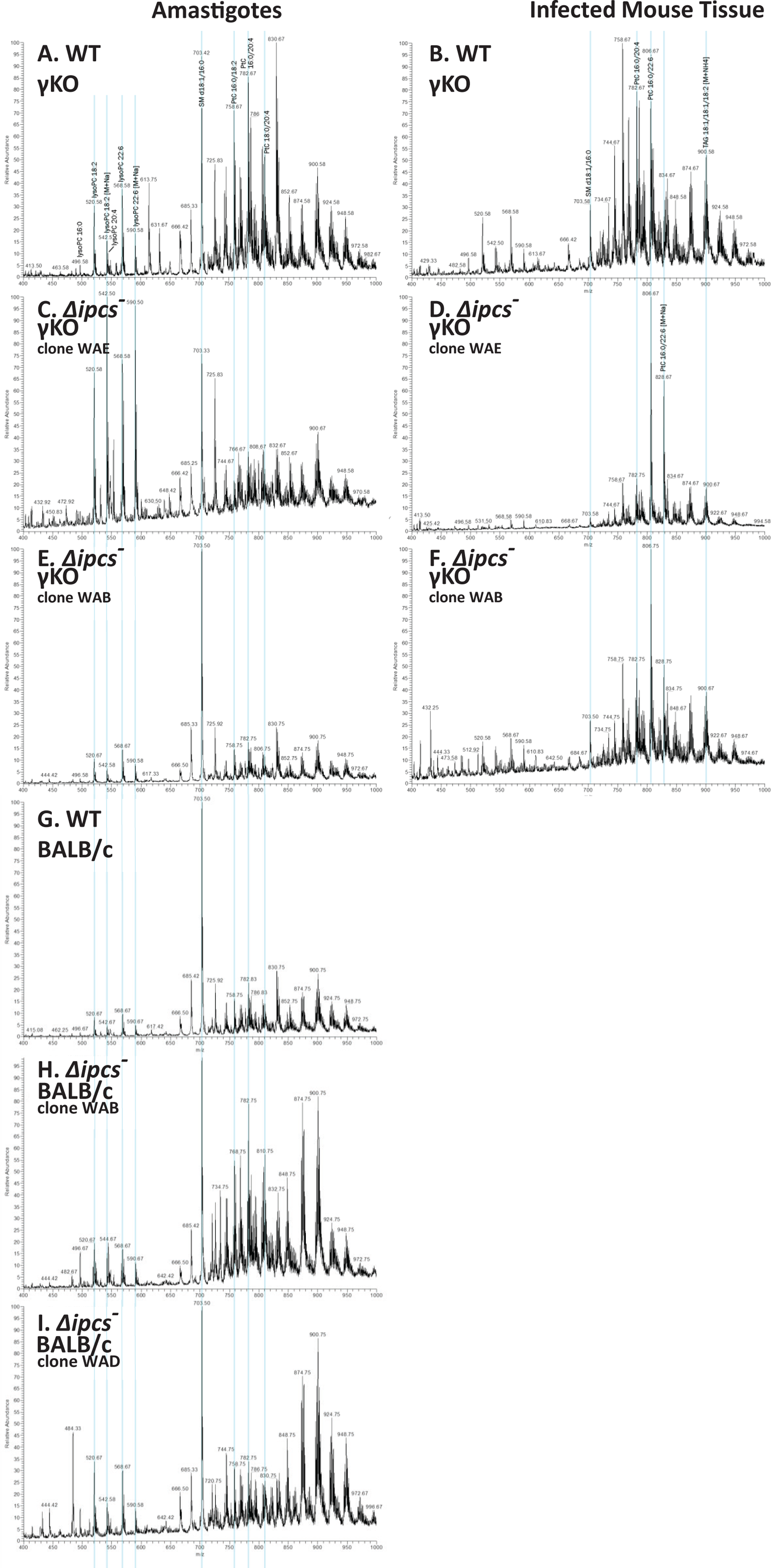
Full scan positive ion mass spectra of the lipid extracts from purified amastigotes or infected mouse tissue. (A), WT amastigotes purified from γKO mice infections. (B), infected lesion tissue from WT infected γKO mice. (C), *Δipcs*^-^ amastigotes (clone WAE) ^purified from^ γKO mice infections. (D), infected lesion tissue from *Δipcs*^-^ (clone WAE) infected γKO mice. (E), *Δipcs*^-^ amastigotes (clone WAB) purified from γKO mice infections. (F), infected lesion tissue from *Δipcs*^-^ (clone WAB) infected γKO mice. (G), WT amastigotes purified from BALB/c mice infections. (H), *Δipcs*^-^ amastigotes (clone WAE) purified from BALB/c mice infections. (I), *Δipcs*^-^ amastigotes (clone WAB) purified from BALB/c mice infections. Major ions for comparison were lined up across the panels. Lipid species abbreviations follow those recommended by the Lipid Maps consortium (www.lipidmaps.org). *Please note that due to the size, Fig. S8 can be found as a separate on-line file*.

**Supplemental Table 1.**
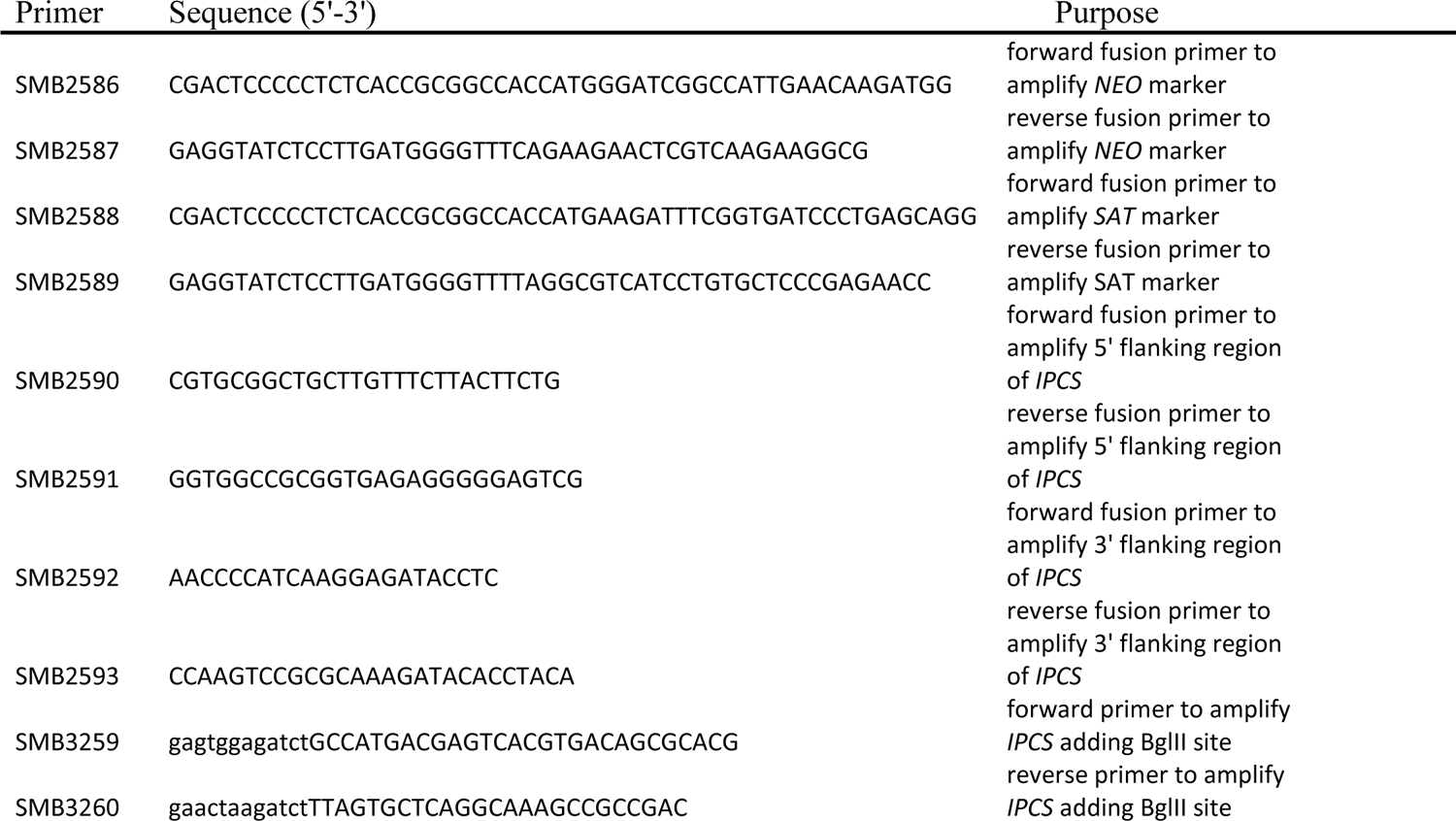
Oligonucleotides used in this work.

**Supplemental Table 2.** Growth of *Δipcs*^-^ *in vitro* is strongly inhibited below pH 5.5 The mean doubling time (hr) and 1 standard deviation (n = 4) of WT, *Δipcs*^-^ or *Δipcs*^-^*/+IPC* with the indicated pH and overlapping buffers is shown.

| Buffer | pH | WT | $\Delta ipcs^-$ | $\Delta ipcs^-/+IPC$ |
| --- | --- | --- | --- | --- |
| Citrate | 5 | 19.7 ± 0.3 | 59.9 ± 18.6* | 20.8 ± 2.1 |
|  | 5.5 | 10.8 ± 0.0 | 15.3 ± 3.7 | 11.3 ± 0.9 |
|  | 6 | 9.5 ± 0.1 | 12.3 ± 1.8 | 10.1 ± 0.4 |
| MES | 5 | 66.8 ± 25.6 | 390.0 ± 119.3* | 35.0 ± 29.1 |
|  | 5.5 | 11.3 ± 1.4 | 23.6 ± 11.8 | 10.4 ± 1.3 |
|  | 6 | 8.8 ± 0.4 | 8.0 ± 4.2 | 9.2 ± 0.5 |
| HEPES | 6.8 | 8.4 ± 0.2 | 10.7 ± 0.1* | 9.3 ± 0.4 |

\* $p < 0.05$ as determined by ANOVA with Tukey's post-hoc analysis; WT vs $\Delta ipcs^-$ .

**Supplemental Table 3.** Levels of C16 and C18 SLs present in several mammalian macrophage sources. The average and range seen in thioglycolate-elicited, bone marrow-derived, and RAW264.7 macrophages is shown. Data were obtained from the Lipid Maps website (www.lipidmaps.org) (1, 2). Units are pM lipid / μg DNA. Shading depicts an arbitrary cutoff for those SLs showing values >1.

| Lipid Maps | Lipid Maps Analyte | Anchor | Average | Range |
| --- | --- | --- | --- | --- |
| ***C16-18 Ceramides*** |  |  |  |  |
| LMSP02010004 | C16 Cer | d18:1/16:0-Cer | 4.23 | (2.48 - 6.82) |
| LMSP02050002 | C16 CerP | d18:1/16:0-Cer-1-P | 0.15 | (0 - 0.42) |
| LMSP02050003 | C16DH CerP | d18:0/16:0-Cer-1-P | 0.00 | (0 - 0.01) |
| LMSP02020001 | C16DH Cer | d18:0/16:0-Cer | 2.58 | (0.63 - 4.25) |
| LMSP02010003 | C18:1 Cer | d18:1/18:1-Cer | 0.04 | (0.03 - 0.06) |
| LMSP02020015 | C18:1DH Cer | d18:0/18:1-Cer | 0.24 | (0.09 - 0.44) |
| LMSP02010006 | C18 Cer | d18:1/18:0-Cer | 0.19 | (0.13 - 0.27) |
| LMSP02050004 | C18 CerP | d18:1/18:0-Cer-1-P | 0.06 | (0 - 0.17) |
| LMSP02020008 | C18DH Cer | d18:0/18:0-Cer | 0.43 | (0.28 - 0.61) |
| ***C16--C18 Glc-Ceramides*** |  |  |  |  |
| LMSP0501AA03 | C16 GlcCer | d18:1/16:0-GlcCer | 6.29 | (2.73 - 12.87) |
| LMSP0501AA04 | C16DH GlcCer | d18:0/16:0-GlcCer | 0.72 | (0.18 - 1.53) |
| LMSP0501AA29 | C18:1DH GlcCer | d18:0/18:1-GlcCer | 0.00 | (0 - 0.01) |
| LMSP0501AA27 | C18:1 GlcCer | d18:1/18:1-GlcCer | 0.01 | (0 - 0.03) |
| LMSP0501AA19 | C18DH GlcCer | d18:0/18:0-GlcCer | 0.06 | (0.01 - 0.11) |
| LMSP0501AA05 | C18 GlcCer | d18:1/18:0-GlcCer | 0.32 | (0.11 - 0.46) |
| ***C16-C18 Sphingomyelins*** |  |  |  |  |
| LMSP03010004 | C16DH Sphingomyelin | d18:0/16:0-SM | 29.34 | (15.98 - 42.56) |
| LMSP03010003 | C16 Sphingomyelin | d18:1/16:0-SM | 191.94 | (88.31 - 314.65) |
| LMSP03010020 | C18DH Sphingomyelin | d18:0/18:0-SM | 1.46 | (0.59 - 2.76) |
| LMSP03010031 | C18:1DH Sphingomyelin | d18:0/18:1-SM | 6.84 | (5.97 - 7.49) |
| LMSP03010029 | C18:1 Sphingomyelin | d18:1/18:1-SM | 0.69 | (0.57 - 0.88) |
| LMSP03010001 | C18 Sphingomyelin | d18:1/18:0-SM | 6.52 | (6.03 - 7.22) |
| ***Sphingoid bases*** |  |  |  |  |
| LMSP01020001 | Sphinganine | d18:0-SB | 0.97 | (0.07 - 2.62) |
| LMSP01050002 | Sphinganine-P | d18:0-SB-1-P | 23.57 | (0.01 - 70.44) |
| LMSP01010001 | Sphingosine | d18:1-SB | 2.06 | (0.76 - 3.12) |
| LMSP01050001 | Sphingosine-P | d18:1-SB-1-P | 0.91 | (0.01 - 2.68) |

## Footnotes

1 Data not shown.

